# In silico codon optimization of the variant antigen-encoding genes of diverse strains of Zika Virus

**DOI:** 10.1101/2023.11.12.566740

**Authors:** Jahnvi Hora, Vanshika Panwar, Indra Mani

## Abstract

With the recent emergence of the coronavirus pandemic, the scare for other viral pandemics is on the rise. Already having caused an epidemic in 2015-16, Zika virus (ZIKV) poses a threat for potential havoc. Thus, there is an urgent need to produce drugs and vaccines for the same as no approved vaccinations exist for the virus as of now. With the optimization of codon usage, the process of vaccine development can be accelerated and efficacy can be ensured. ZIKV genome has 3 structural genes (envelope, capsid and membrane which is obtained from pre membrane) and 7 nonstructural genes (NS1, NS2A, NS2B, NS3, NS4A, NS4B and NS5). We have used these 10 genes in 5 different strains of ZIKV for in silico DNA optimization in *Escherichia coli.* The mean CAI, GC% and AT% of wild-type DNA and optimized DNA of each were compared. It was observed that the CAI and GC% had increased in optimized DNA as compared to wild-type while the opposite was seen for AT%. The results show that codon optimization helps in efficient expression of proteins in the host. These could be used in the development of biotherapeutics. The ideal genes to be overexpressed in development of biotherapeutics are the membrane precursor (prM) and envelope (E) genes as the results have shown.

## 1. Introduction

Zika Virus (ZIKV), initially limited to scattered and irregular cases across Africa and Asia, gained sudden attention during the outbreak in the 21^st^ century. Its emergence in Brazil (2015) led to a sudden spread throughout the Americas. The virus belongs to the genus Flavivirus and the family Flaviviridae (Huang et al., 2014). The family also includes dengue virus (DENV), yellow fever virus (YFV) and West Nile virus (WNV). It is an arthropod-borne virus, transmitted to humans by the bite of infected female Aedes mosquito (*Ae. aegypti* and *Ae. albopictus*) (Dick et al., 1952). It is usually responsible for causing asymptomatic infection but neurological disorders like microcephaly, a condition in which the fatal brain fails to develop properly or severe neurological complications like Guillain-Barré syndrome as well as congenital Zika syndrome may arise (Alcantra et al., 2014; Ansar et al., 2015; Pierson et al., 2018). It was first isolated from a feverish rhesus macaque monkey in 1947 in the Zika Forest of Uganda, later, it was identified in the Aedes africanus mosquitoes in the same forest (Dick et al., 1952). In Africa, it has been isolated in many other genera of mosquitoes like; Aedes, Anopheles, and Mansonia. It has been isolated from monkeys, but its antibodies have been found in ducks, cows, orangutans, bats, sheep, goats, horses, and rodents (Olson et al., 1983). The Zika virus particles are small and spherical (around 50nm in diameter), and have a ‘golf ball’ structure with single stranded RNA genome. The virions have an envelope and a nucleocapsid. Surface protein E (envelope) is parallel to viral membrane which gives it a smooth surface. The surface consists of 180 copies of E and M (membrane) proteins embedded in the viral membrane in icosahedral symmetry (Sirohi et al., 2017). The electron dense core is 30 nm in diameter. ZIKV is structurally and functionally similar to other flavivirus pathogens like Japanese encephalitis, dengue and West Nile viruses. African and Asian are the two major lineages of ZIKV (Beaver et al., 2018). Almost all outbreaks and birth defects have occurred due to the Asian lineage, although laboratory studies suggest that the African lineage shows higher transmissibility and fetal pathogenicity, thus becoming a potential epidemic scare (Aubry et al., 2021).

The genome of ZIKV consists of a positive sense, single stranded RNA and a size of almost 10.8 kb and a single open reading frame (ORF) flanked with 5′- and 3′-untranslated regions (UTRs). The translated RNA gives a single polyprotein, 3423 amino acids long. This encodes 3 structural proteins- envelope (E); capsid (C); and membrane (M), generated from its precursor pre membrane (prM)- along with 7 nonstructural proteins (NS1, NS2A, NS2B, NS3, NS4A, NS4B, NS5) (Wang et al., 2017; Pielnaa et al., 2020). The structural proteins are responsible for assembly of the virus particle and its pathogenicity, while the non-structural proteins help in replication and packaging of the genome and disrupting host pathways to aid the virus. The capsid protein is responsible for the icosahedral viral capsid, surrounded by a spherical lipid bilayer membrane derived from the host. The E protein is involved in binding of host cells and membrane fusion and is the major virion surface protein. The prM protein is cleaved into the M protein and pr peptide by a furin-like protease. prM participates in E protein folding. The M and E proteins get displayed on the surface and have transmembrane helices which assist in anchoring them in the outer membrane (Wang et al., 2017). All 7 non-structural proteins play a role in host innate immunity hostility and form the replicative complex. NS1 protein of ZIKV is like other Flaviviruses. It acts as a potential antiviral target and is also involved in immune invasion. It plays a role in viral replication along with NS4A. NS3 is essential in viral replication and processing of polyprotein. NS2A, NS2B, NS4A and NS4B may be membrane associated as they are hydrophobic. NS5 is involved in; RNA dependent RNA polymerase activity and RNA capping. It also suppresses IFN signalling (Wang et al., 2017). The E protein structure exposes differences between flaviviruses and the regions surrounding the glycosylated area varies among different flaviviruses which may be an important determinant for specificity of antibody. For ZIKV, the Asn154 site is glycosylated in many strains, this is also found in DENV E protein (Wang et al., 2017).

The virus is primarily spread by the bites of infected female *Ae. aegypti* and *Ae. albopictus* mosquitoes which have been known to cause Dengue and Chikungunya as well. The mosquitoes get infected once they bite a person with ZIKV during the first week of infection (the virus can be found in the blood during that time). They may bite during the day or night. The virus may also be transmitted from mother to the child. During pregnancy, it may result in microcephaly and other brain defects. The virus has also been found in breast milk however infection via that route has not been confirmed yet. The virus has also been known to be transmitted via sexual intercourse and blood transfusions. Although majorly asymptomatic, symptomatic cases present mild cases mostly. The symptoms are manifested in the form of rash, fever, arthralgia, conjunctivitis, and myalgia. However, neurological (microcephaly in newborn babies) and autoimmune complications (Guillain-Barré syndrome) have also been observed in ZIKV patients (Musso et al., 2019). RNA viruses mutate easily, but it is not necessary that mutations make the virus virulent than before or its host more efficient. However, it has been observed by whole genome comparative analysis of earlier and presently endemic ZIKV strains that there have been several mutations that might have contributed in turning Zika virus lethal (Zhu et al., 2016). The virus from the Asian lineage (that has caused the epidemics) has a mutation in the NS1 gene that surfaced after 2012 and is responsible for enhanced mosquito infection. Results have also shown that accumulation of mutations increases evasion of the virus by the immune response and raises chances of potential epidemics (Xia et al., 2018). As the world is already suffering from a pandemic, it becomes the need of the hour to study more about other viruses that may pose threats as a potential pandemic and develop efficient vaccinations for the same.

Currently, there are no approved vaccines for Zika virus. The alarming increase in the number of Zika virus cases has led to a sudden urge to brainstorm ideas to develop vaccines (Shan et al., 2018). As there already exist vaccines for the other four flaviviruses- Yellow Fever Virus (live-attenuated), Tick-borne encephalitis (inactivated), Japanese encephalitis virus (both inactivated and live-attenuated), and dengue virus (chimeric live-attenuated)- it is practical to have a safe and potent Zika virus vaccine. Lessons learned from the development of these vaccines can guide the development of a Zika virus vaccine. **Table 1** summarizes the progress in development of the Zika virus vaccine.

**Table 1.**
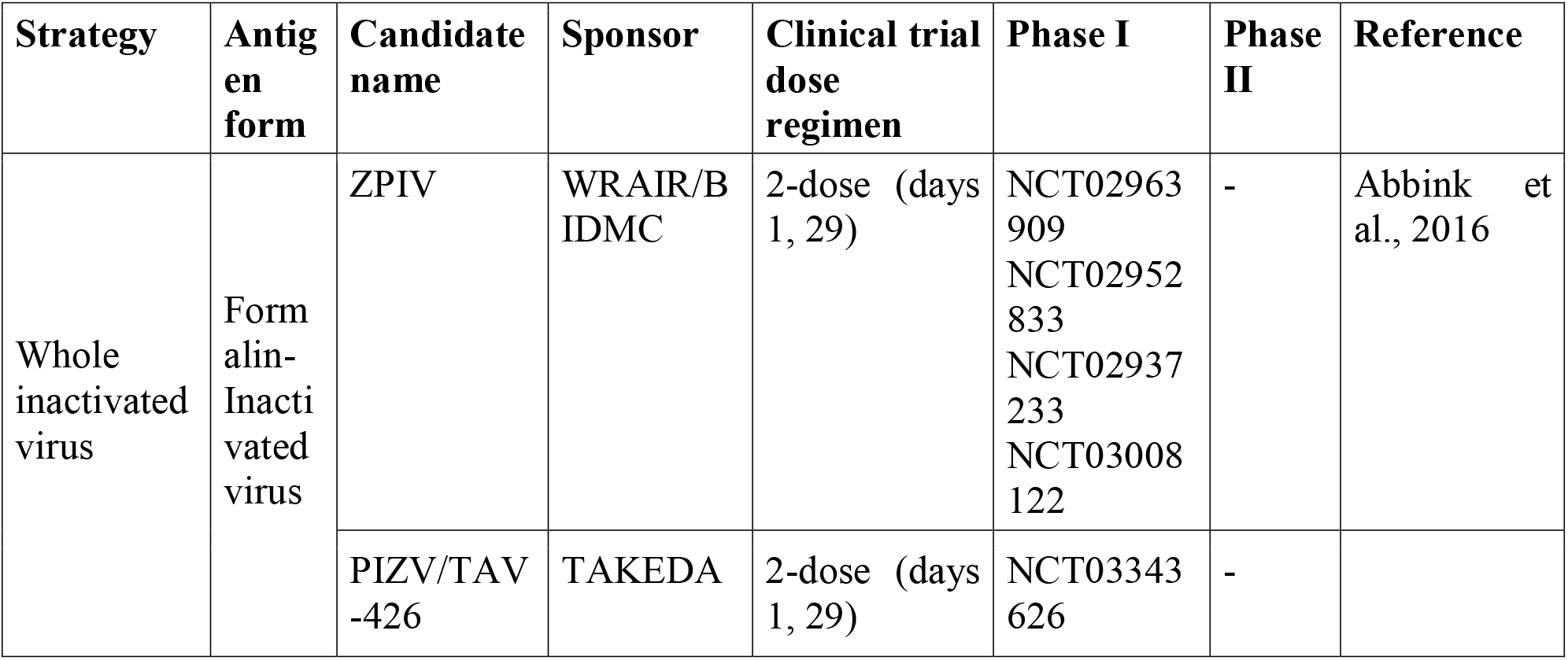
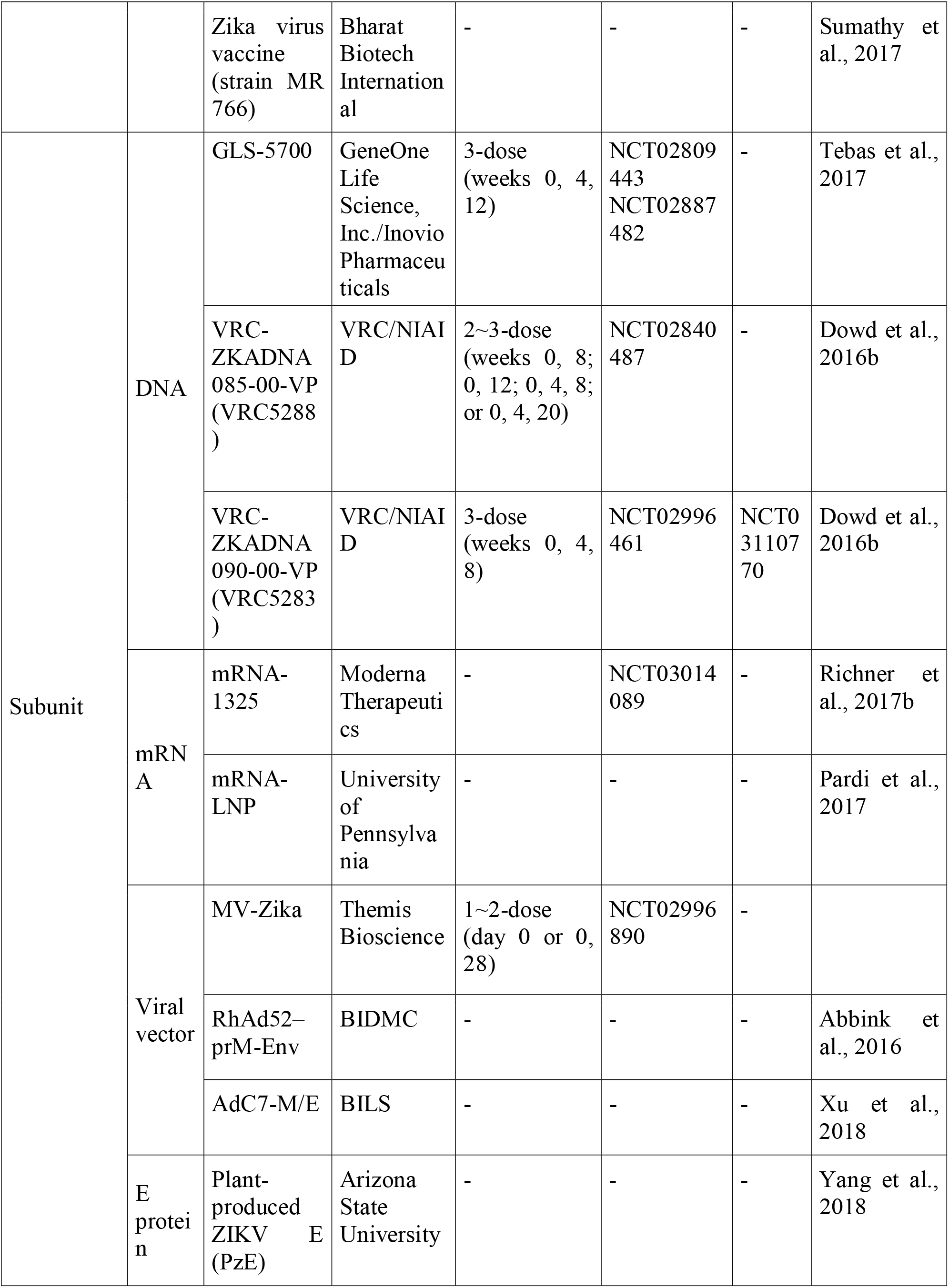
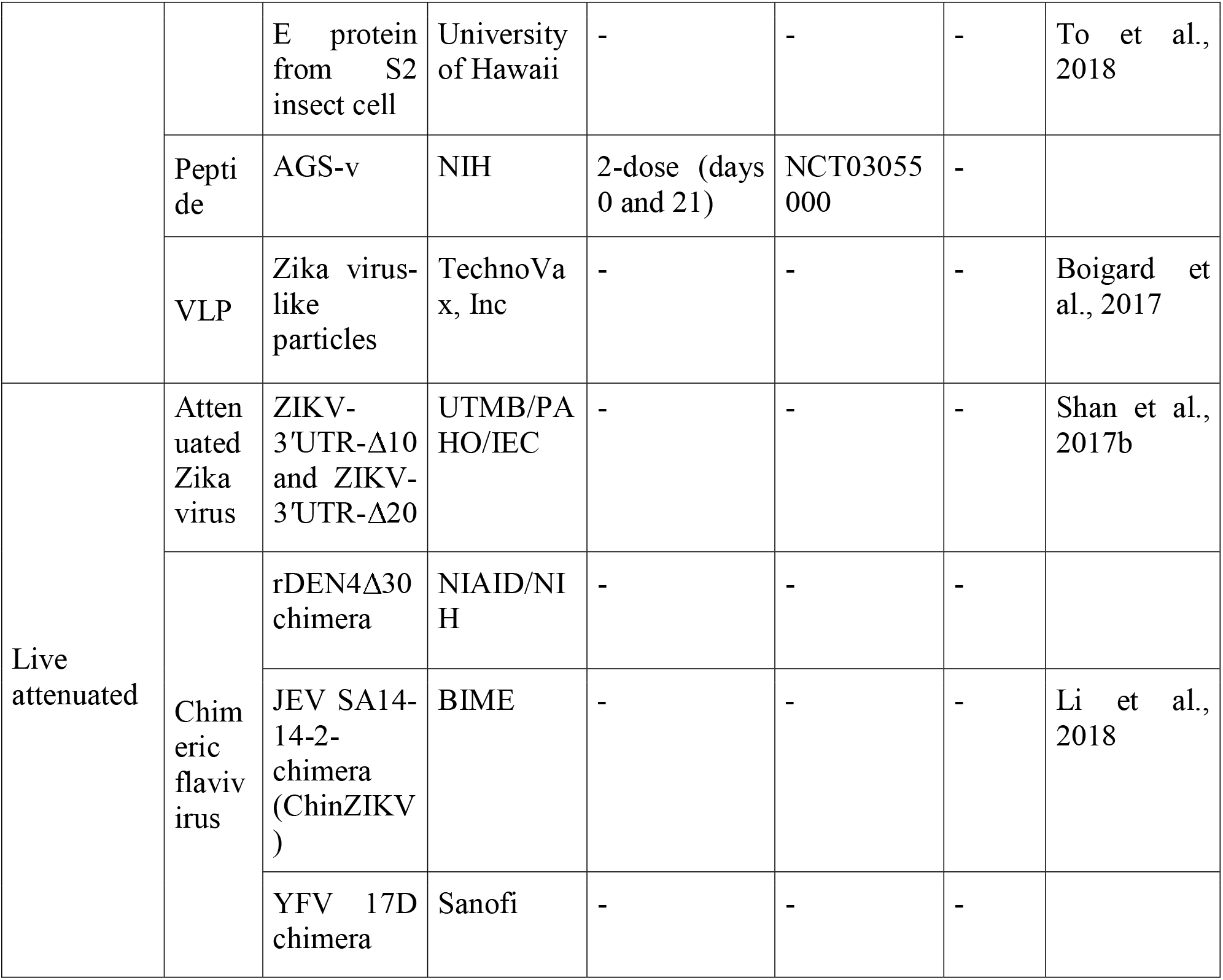
Zika Vaccines in Development.

The vaccine candidates belong to three distinct categories: (i) inactivated vaccine, (ii) subunit vaccine, and (iii) live-attenuated vaccine (Shan et al., 2018). Some vaccine candidates have been close to success, for example, the E ZIKV antigen vaccine (Kim et al., 2018), vaccine using self-amplifying messenger RNA (Luisi et al., 2020), virus-like particle (VLP)-based approach to develop a vaccine and a microneutralization assay for ZIKV (Garg et al., 2017) etc. However, these have not been approved. Studies show that in DNA vaccines, the immunogenicity increases as the expression gets maximized in the host. Codon bias can thus be used to enhance the optimization of the protein product in the vaccine for the virus which generates effective immunity (Hemert et al., 2016). No specific and approved drugs exist for the virus either. General medicines for fever, rashes, and fluids to prevent dehydration are given. The ongoing search for antivirals against the virus has approaches targeting the various steps of the replication cycle. The antivirals could be directly acting or host targeting (Baz et al., 2019). As the world is already suffering from a pandemic, it becomes the need of the hour to study more about other viruses that may pose threats as a potential pandemic and develop efficient vaccinations for the same and this is what the study aims to accomplish.

As for antivirals, no specific drugs have been approved for treating ZIKV. For controlling pain and fever acetaminophen is used, for pruritic rashes antihistamines are used and to prevent dehydration fluids are given. As there is an increased risk of haemorrhagic syndrome reported with other flaviviral infections along with risk of Reye’s syndrome in children and teenagers, acetylsalicylic acid, and non-steroidal anti-inflammatory drugs (NSAIDs) are also suggested. Currently, the search for the antiviral options against ZIKV has varied approaches, mostly targeting the different steps of the replication cycle. Depending on mode of action, antivirals are classified as: direct-acting antivirals and host-targeting antivirals (Baz et al., 2019). Optimization of a codon is done in order to refine codon composition of a gene without altering the amino acid sequence. This can be done because of degeneracy of codons (Mauro et al., 2014). The process of codon optimization is regularly used to increase efficiency of translation of a gene of interest by transforming the DNA sequence of one species to that of another; human to bacteria etc. (Graf et al., 2009). Codon usage bias is the occurrence where some codons are used more frequently than other codons (synonymous) during translation. This varies among species. It is an important factor to determine gene expression and cellular function and affects processes like RNA processing, protein translation and folding. It shows the origin, patterns of mutation and evolution of genes or species (Parvathy et al., 2021). Codon usage variation in ZIKV is affected by natural selection such as aromaticity, hydropathicity host, geography etc. and codon usage bias is low. Studying codon usage patterns allows us to study the molecular evolution patterns and also forms the basis of development of therapeutics and vaccines (Tao et al., 2020). The aim of the study was to optimize the codon for over-expression of different zika virus genes in *E. coli* to develop the vaccine and produce it in adequate amounts. We propose that the studied genes might be useful for increasing protein expression levels to ensure the efficient production of vaccines.

## 2. Methods

In the present study, we optimized the codons for over-expression of different zika virus genes in *E. coli* to develop the vaccine and produce it in adequate amounts.

### 2.1 Collection of sequences

From NCBI-GenBank site (http://www.ncbi.nlm.nih.gov), nucleotide sequences of different genes of Zika virus (Accession numbers: MW015936.1, MT078739.1, KY785429.1, OK054385.1, MW122391.1) were retrieved (Benson et al., 1994).

### 2.2 Codon optimization and analysis

For optimization and calculation of codon adaptation index (CAI), G+C and A+T content of the retrieved DNA sequences with reference to *E. coli* K-12 MG1655, an optimizer is used which is a tool for (http://genomes.urv.es/OPTIMIZER/) (Puigbo et al., 2007) anticipating and optimizing gene expression levels in a heterologous expression host. *E. coli* K-12 MG1655 is used as it is a common host for heterologous gene expression. CAI was calculated for all 5 strains.

### 2.3 Statistical analysis

Using GraphPad Prism (version 8.1) software, the mean, range and standard deviation can be calculated for statistical analysis. With the values, a graph can be plotted and CAI of wild-type is to be compared with the optimized sequences amongst various strains of the Zika virus. The comparison between the CAI, GC, and AT of all 10 genes of Zika virus was drawn using the Mann Whitney test. A two-tailed probability p<0.05 was considered for statistical significance.

### 2.4 Nucleotide sequence alignment

Clustal omega was used to carry out nucleotide sequence alignment between wild-type and optimized sequences of all 10 genes of strains MW015936.1, MT078739.1, KY785429.1, OK054385.1, MW122391.1.

## 3. Results

As of now there are no approved vaccines or drugs (antivirals) for zika virus. Thus, the need to develop effective vaccines is dire in order to prevent another epidemic or pandemic. Through this study, codon optimization can bring about the efficiency of translation of desired proteins in the host. The codon usage of various genes of Zika virus (Capsid, envelope, precursor membrane, NS1, NS2A, NS2B, NS3, NS4A, NS4B and NS5) have been recorded in the tables below. The codons were optimized in the host *E. coli* K-12 MG1655. It was observed that the CAI of optimized genes was 1 in all cases which is the maximum value and implies that the gene always uses the most frequently used synonymous codons in the reference set.

The range of the CAI, GC% and AT% frequencies for the 5 strains of wild-type capsid protein are from 0.191 to 0.217, 47.8% to 49.2% and 50.8% to 52.2% respectively and a mean (±SD) of 0.201 ± 0.01, 48.3 ± 0.6 and 51.6 ± 0.6 respectively. The frequencies for the same in optimized DNA range from 1 to 1, 47.9 to 51.6 and 48.4 to 50.3 with a mean (±SD) of 1 ± 0, 50.48 ± 0.94 and 49.5 ± 0.94 respectively. When the average CAI, GC% and AT% of wild-type and optimized DNA were compared, significant changes were observed. The mean CAI and GC% in optimized DNA were 4.97 (397.51%) and 1.04 (4.51%) fold higher respectively than their wild-type counterparts. But the mean AT% in optimized DNA was decreased by 4.24% from wild-type (Table 2).

**Table 2.**
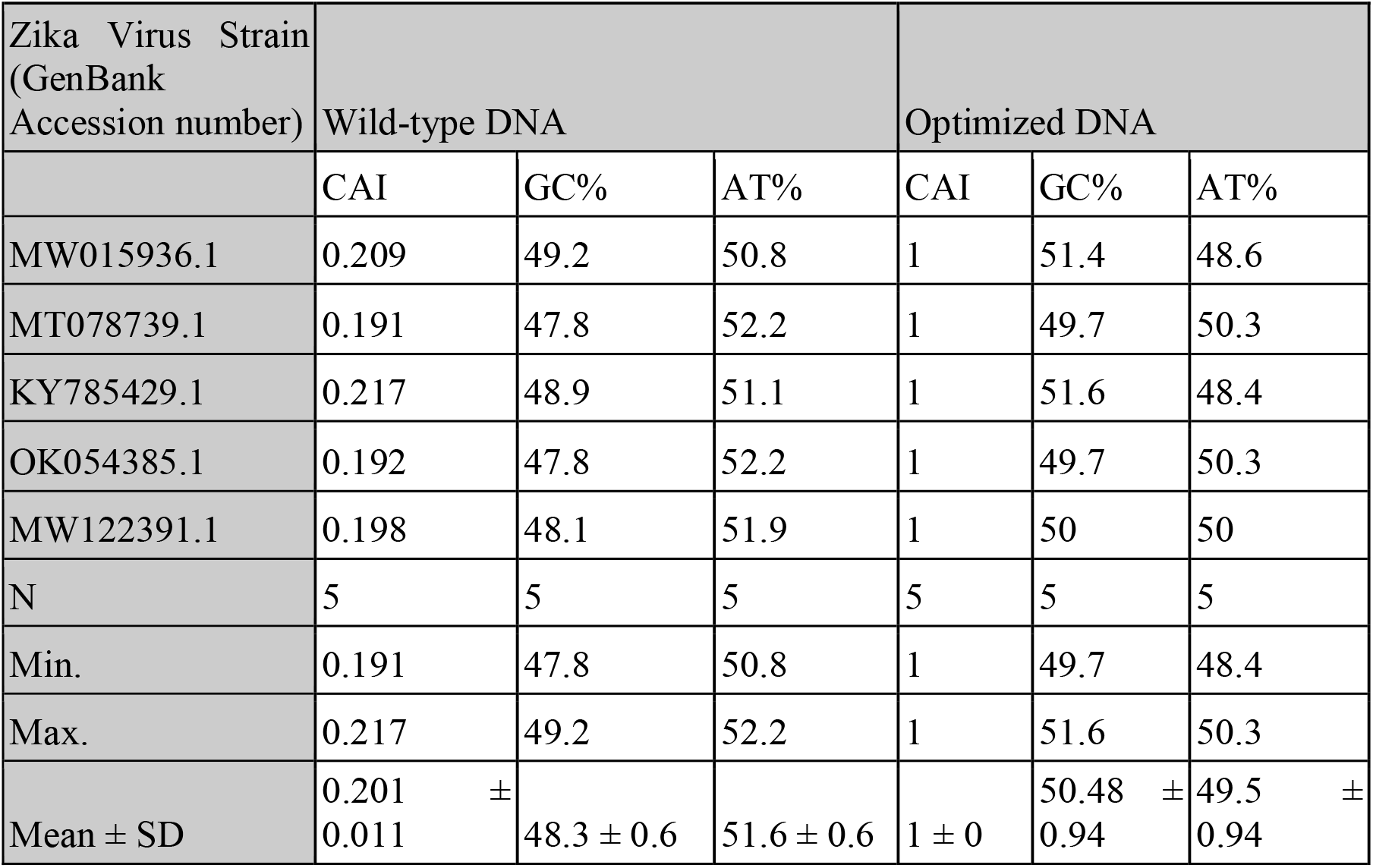
Capsid protein (C) gene of Zika virus expression level in *E. coli* of the wild-type and optimized DNA sequences.

The alignment of the wild and optimized sequences for the capsid gene of different strains are also shown in supplementary figure S1. A graph showing the CAI values on the X-axis and the strains on the Y-axis is given in **Figure 1**.

**Figure 1.**
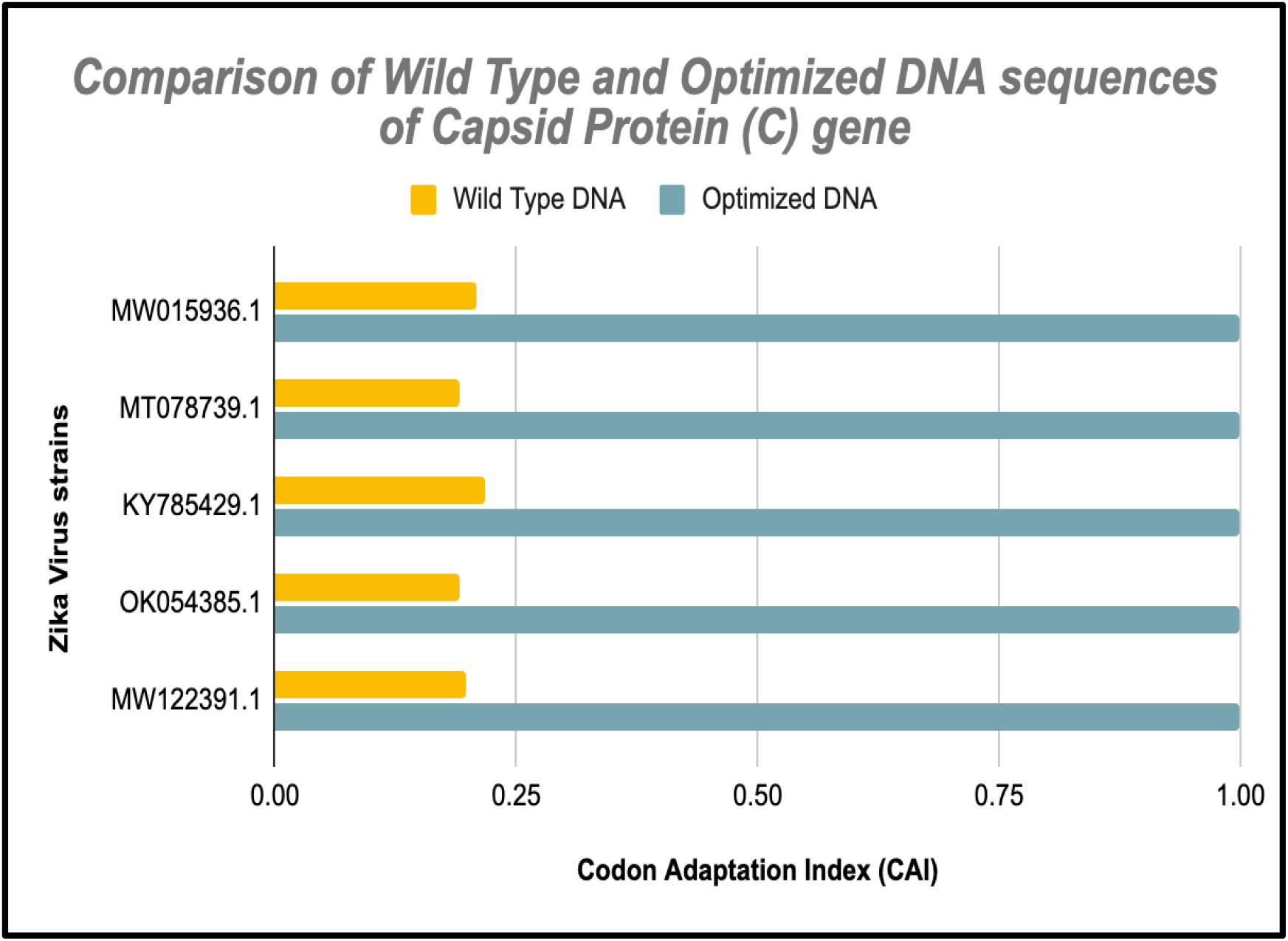
Comparison of the wild-type and optimized DNA sequences of nucleocapsid gene in different strains of ZIKA virus.

Likewise, the CAI, GC% and AT% frequencies of membrane precursor protein (prM) in wild-type strains range from 0.260 to 0.296, 48% to 49.2% and 50.8% to 52% respectively with an average (±SD) of 0.268(±0.0155), 48.8(±0.4) and 51.1(±0.4) respectively (Table 3). Their respective frequencies in optimized DNA range from 1 to 1, 55.2% to 55.8% and 44.2% to 44.8% and mean (±SD) of 1(±0), 55.6(±0.2) and 44.4(±0.2). The mean CAI and GC in optimized DNA were 3.73(273.1%) and 1.13(13.9%) fold higher than respective wild-type strains. But, the mean of AT content in optimized DNA was decreased by 13.11% compared to wild-type.

**Table 3.**
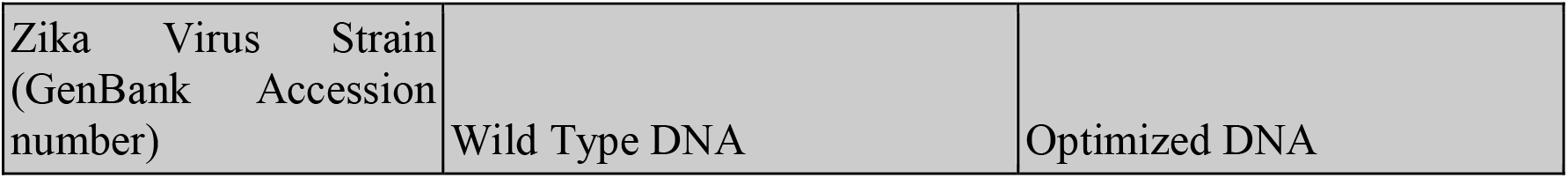
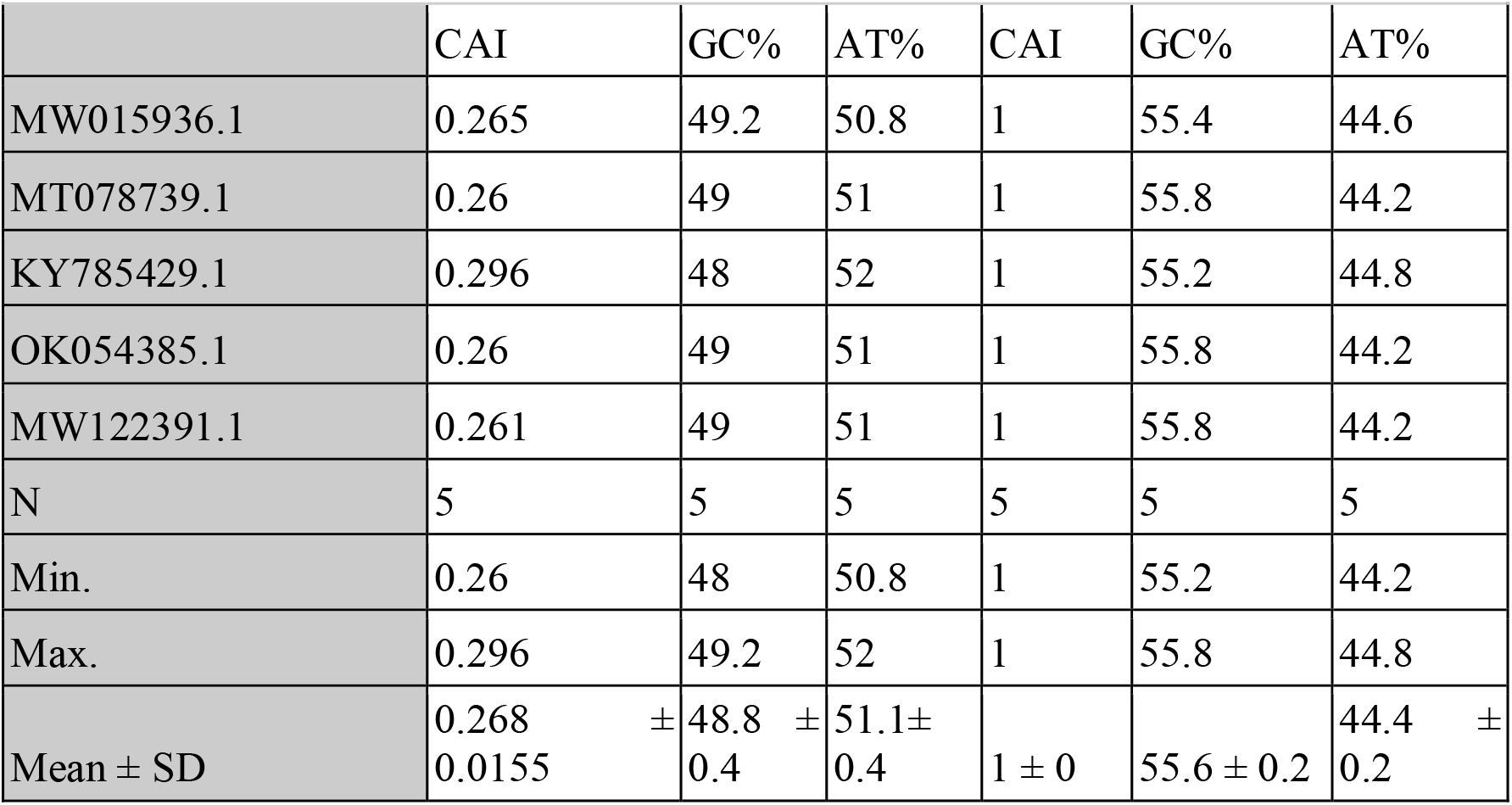
Membrane precursor (prM) gene of Zika virus expression level in *E. coli* of the wild-type and optimized DNA sequences.

| Zika Virus Strain<br>(GenBank Accession<br>number) | Wild Type DNA | Optimized DNA |
| --- | --- | --- |

|  | CAI | GC% | AT% | CAI | GC% | AT% |
| --- | --- | --- | --- | --- | --- | --- |
| MW015936.1 | 0.265 | 49.2 | 50.8 | 1 | 55.4 | 44.6 |
| MT078739.1 | 0.26 | 49 | 51 | 1 | 55.8 | 44.2 |
| KY785429.1 | 0.296 | 48 | 52 | 1 | 55.2 | 44.8 |
| OK054385.1 | 0.26 | 49 | 51 | 1 | 55.8 | 44.2 |
| MW122391.1 | 0.261 | 49 | 51 | 1 | 55.8 | 44.2 |
| N | 5 | 5 | 5 | 5 | 5 | 5 |
| Min. | 0.26 | 48 | 50.8 | 1 | 55.2 | 44.2 |
| Max. | 0.296 | 49.2 | 52 | 1 | 55.8 | 44.8 |
| Mean ± SD | 0.268 ± 0.0155 | 48.8 ± 0.4 | 51.1 ± 0.4 | 1 ± 0 | 55.6 ± 0.2 | 44.4 ± 0.2 |

A graph was plotted and the CAI value was shown at X-axis, while the number of studied strains on Y-axis (Figure 2). The alignment of the wild and optimized sequences for the prM gene of different strains are also shown in supplementary figure S2.

**Figure 2.**
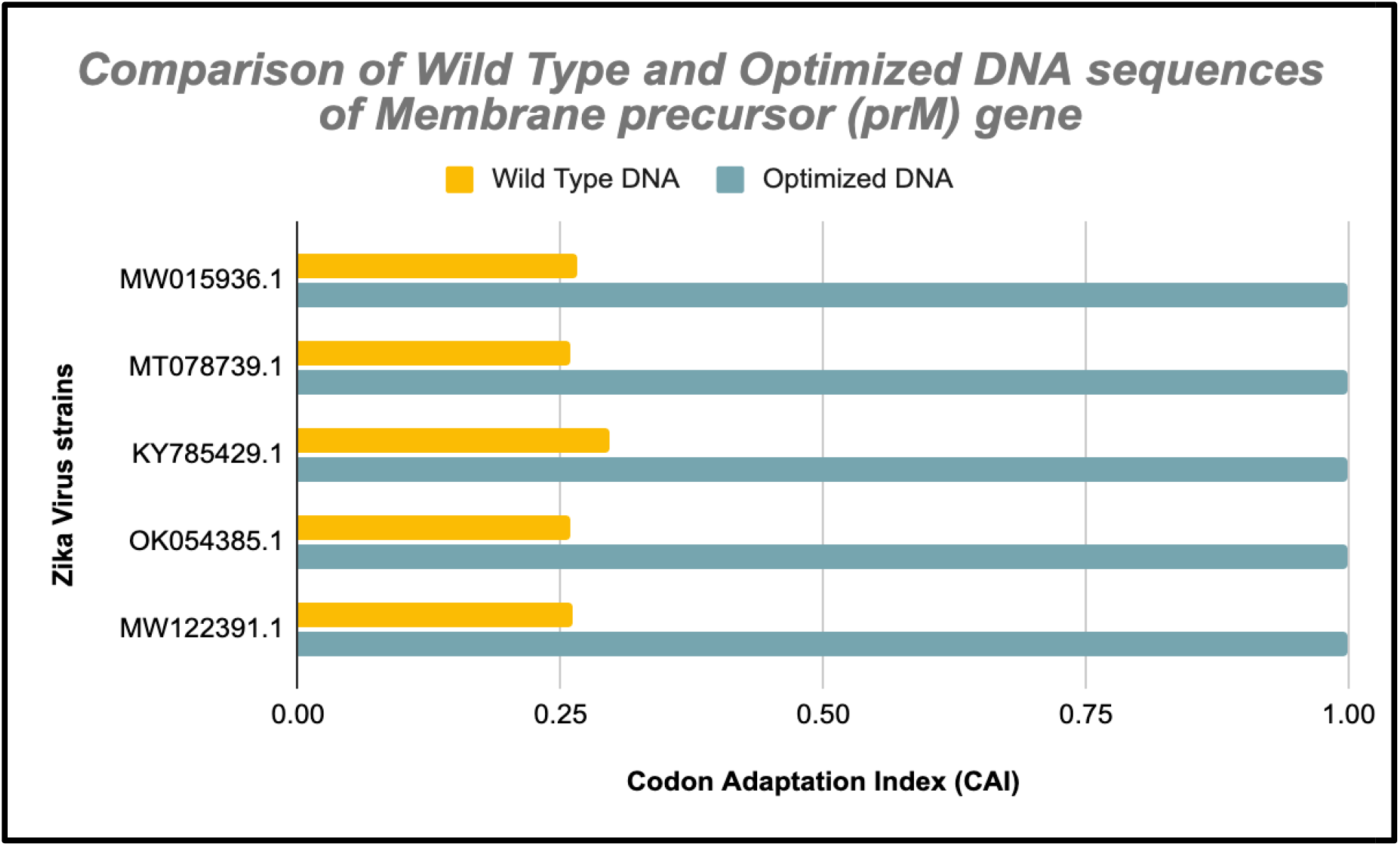
Comparison of the wild-type and optimized DNA sequences of membrane precursor (prM) gene in different strains of ZIKA virus.

The CAI, GC% and AT% frequencies of envelope (E) gene in wild-type DNA range from 0.294 to 0.301, 49.3% to 49.8% and 50.2% to 50.7% respectively and an average (±SD) of 0.297(±0.002), 49.6(±0.1), 50.4(±0.1) respectively. The respective frequencies of these in optimized DNA range from 1 to 1, 52.2% to 54.3% and 45.7% to 47.8% with an average (±SD) of 1(±0), 53.8(±0.9), 46.1(±0.9) (Table 4). The mean CAI and GC in optimized DNA were found to be 3.36(236.7%) and 1.08(8.46%) fold higher than respective wild-type values. However, the mean of AT content in optimized DNA was reduced by 8.66% compared to wild-type.

**Table 4.** Envelope (E) gene of Zika virus expression level in *E. coli* of the wild-type and optimized DNA sequences.

| Zika Virus Strain (GenBank Accession number) | Wild-type DNA |  |  | Optimized DNA |  |  |
| --- | --- | --- | --- | --- | --- | --- |
|  | CAI | GC% | AT% | CAI | GC% | AT% |
| MW015936.1 | 0.297 | 49.6 | 50.4 | 1 | 54.3 | 45.7 |
| MT078739.1 | 0.297 | 49.7 | 50.3 | 1 | 54.3 | 45.7 |
| KY785429.1 | 0.301 | 49.3 | 50.7 | 1 | 52.2 | 47.8 |
| OK054385.1 | 0.294 | 49.8 | 50.2 | 1 | 54.3 | 45.7 |
| MW122391.1 | 0.296 | 49.6 | 50.4 | 1 | 54.3 | 45.7 |
| N | 5 | 5 | 5 | 5 | 5 | 5 |
| Min. | 0.294 | 49.3 | 50.2 | 1 | 52.2 | 45.7 |
| Max. | 0.301 | 49.8 | 50.7 | 1 | 54.3 | 47.8 |
| Mean ± SD | 0.297 ± 0.002 | 49.6 ± 0.1 | 50.4 ± 0.1 | 1 ± 0 | 53.8 ± 0.9 | 46.1 ± 0.9 |

A graph was plotted and the CAI value was shown at X-axis, while the number of studied strains on Y-axis (Figure 3). The alignment of the wild and optimized sequences for the envelope gene of different strains are also shown in supplementary figure S3.

**Figure 3.**
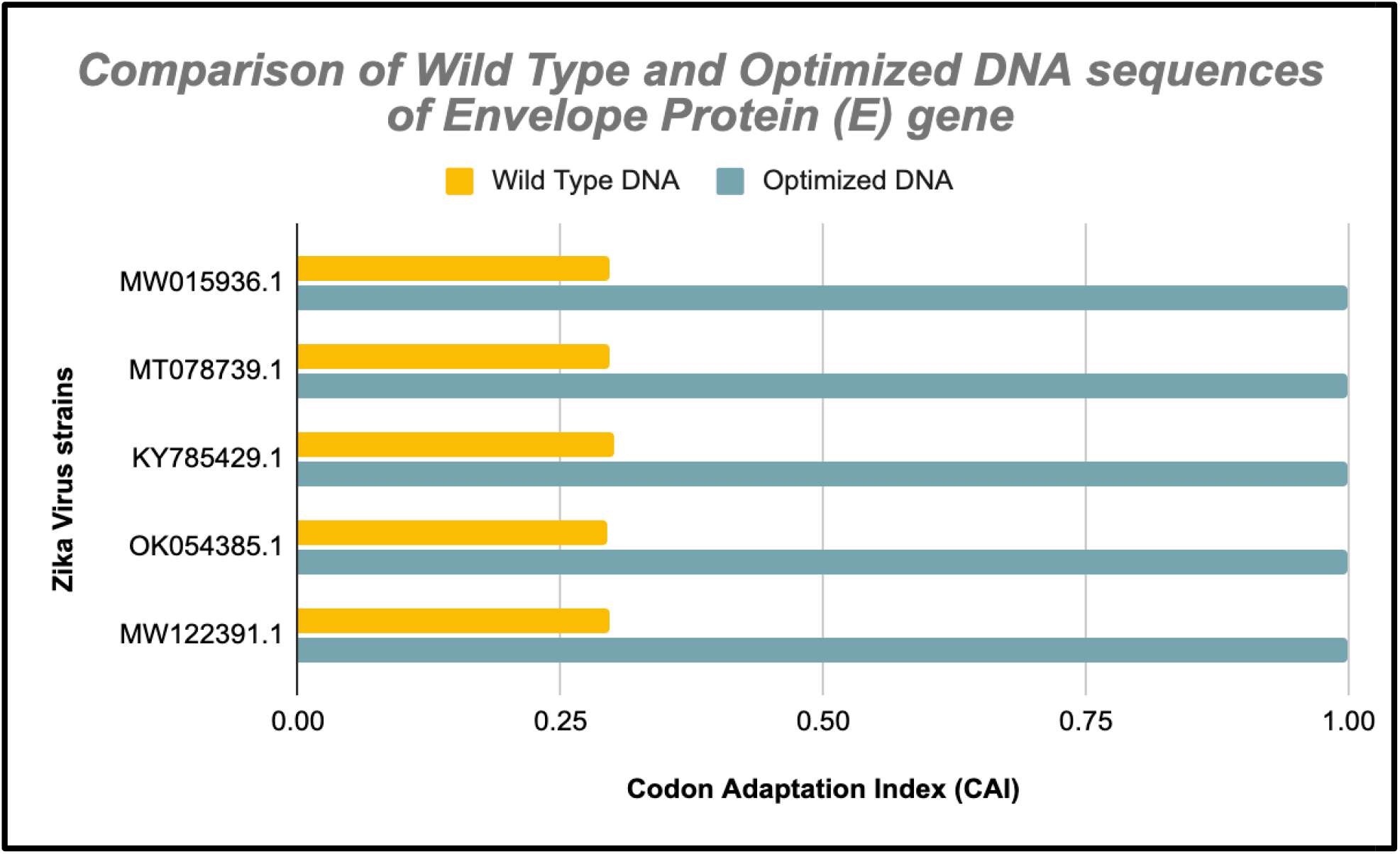
Comparison of the wild and optimized DNA sequences of envelope (E) gene in different strains of ZIKA virus.

Further, the CAI, GC% and AT% frequencies of non-structural protein 1 (NS1) in wild-type strains range from 0.255 to 0.260, 50.2 to 50.5 and 49.5 to 49.8 respectively with an average (±SD) of 0.256(±0.001), 50.3(±0.1) and 49.6(±0.1) respectively. The respective frequencies of these in optimized DNA range from 1 to 1, 52.7 to 52.9 and 47.1 to 47.3 with an average (±SD) of 1(±0), 52.9(±0.08) and 47.2(±0.08) (Table 5). The mean CAI and GC% in optimized DNA were found to be 3.9(290.6%) and 1.05(5.16%) fold higher than respective wild-type strains. But, the mean of AT content in optimized DNA was decreased by 4.83% compared to wild-type.

**Table 5.** non-structural protein 1 (NS1) gene of Zika virus expression level in *E. coli* of the wild-type and optimized DNA sequences.

| Zika Virus Strain<br>(GenBank<br>Accession<br>number) | Wild-type DNA |  |  | Optimized DNA |  |  |
| --- | --- | --- | --- | --- | --- | --- |
|  | CAI | GC% | AT% | CAI | GC% | AT% |
| MW015936.1 | 0.256 | 50.5 | 49.5 | 1 | 52.9 | 47.1 |
| MT078739.1 | 0.26 | 50.3 | 49.7 | 1 | 52.8 | 47.2 |
| KY785429.1 | 0.256 | 50.3 | 49.7 | 1 | 52.7 | 47.3 |
| OK054385.1 | 0.256 | 50.3 | 49.7 | 1 | 52.8 | 47.2 |
| MW122391.1 | 0.255 | 50.2 | 49.8 | 1 | 52.7 | 47.3 |
| N | 5 | 5 | 5 | 5 | 5 | 5 |
| Min. | 0.255 | 50.2 | 49.5 | 1 | 52.7 | 47.1 |
| Max. | 0.26 | 50.5 | 49.8 | 1 | 52.9 | 47.3 |
| Mean ± SD | $0.256 \pm 0.001$ | $50.3 \pm 0.1$ | $49.6 \pm 0.1$ | $1 \pm 0$ | $52.9 \pm 0.08$ | $47.2 \pm 0.08$ |

A graph was plotted and the CAI value was shown at X-axis, while the number of studied strains on Y-axis (Figure 4). The alignment of the wild and optimized sequences for the NS1 gene of different strains are also shown in supplementary figure S4.

**Figure 4.**
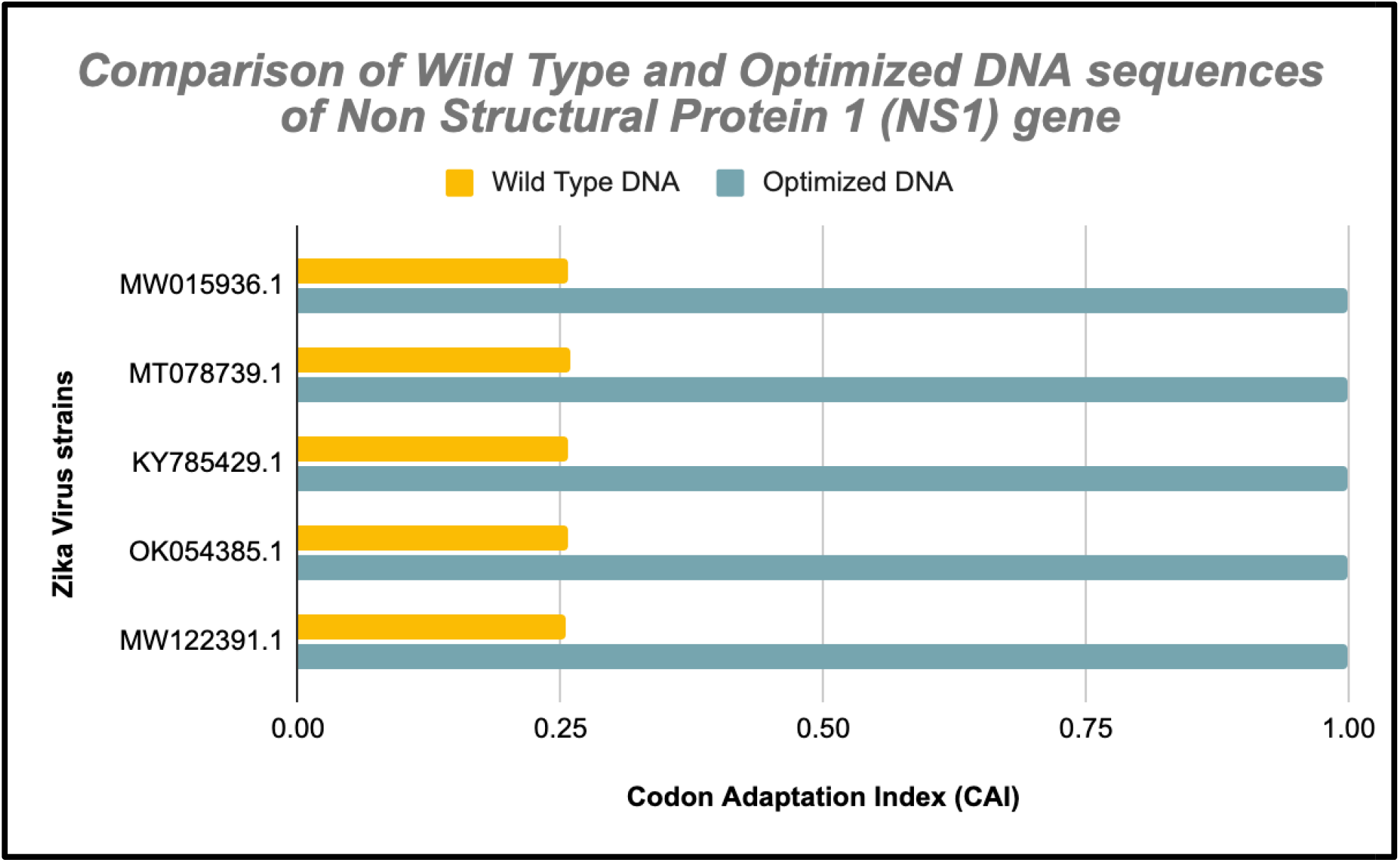
Comparison of the wild and optimized DNA sequences of non-structural protein 1 (NS1) gene in different strains of ZIKA virus.

Next, the CAI, GC% and AT% frequencies of non-structural protein 2A (NS2A) in wild-type strains range from 0.278 to 0.291, 52.2 to 53.4 and 46.6 to 47.8 respectively with an average (±SD) of 0.287(±0.005), 52.7(±0.4) and 47.3(±0.4) respectively; while these values in optimized DNA range from 1 to 1, 57.4 to 57.8 and 42.2 to 42.6 with an average (±SD) of 1(±0), 57.54(±0.15) and 43.46(±0.15) (Table 6). The mean CAI and GC in optimized DNA were found to be 3.48(258.4%) and 1.09(9.1%) fold higher than respective wild-type strains. However, the mean of AT content in optimized DNA was decreased by 8.11% compared to wild-type.

**Table 6.** Non-structural protein 2A (NS2A) gene of Zika virus expression level in *E. coli* of the wild-type and optimized DNA sequences.

| Zika Virus Strain<br>(GenBank<br>Accession number) | Wild-type DNA |  |  | Optimized DNA |  |  |
| --- | --- | --- | --- | --- | --- | --- |
|  | CAI | GC% | AT% | CAI | GC% | AT% |
| MW015936.1 | 0.288 | 53.4 | 46.6 | 1 | 57.8 | 42.2 |
| MT078739.1 | 0.29 | 52.5 | 47.5 | 1 | 57.4 | 42.6 |
| KY785429.1 | 0.291 | 52.7 | 47.3 | 1 | 57.5 | 42.5 |
| OK054385.1 | 0.278 | 52.2 | 47.8 | 1 | 57.5 | 42.5 |
| MW122391.1 | 0.288 | 52.7 | 47.3 | 1 | 57.5 | 42.5 |
| N | 5 | 5 | 5 | 5 | 5 | 5 |
| Min. | 0.278 | 52.2 | 46.6 | 1 | 57.4 | 42.2 |
| Max. | 0.291 | 53.4 | 47.8 | 1 | 57.8 | 42.6 |
| Mean ± SD | 0.287 ± 0.005 | 52.7 ± 0.4 | 47.3 ± 0.4 | 1 ± 0 | 57.54 ± 0.15 | 43.46 ± 0.15 |

A graph was plotted and the CAI value was shown at X-axis, while the number of studied strains on Y-axis (Figure 5). The alignment of the wild and optimized sequences for the NS2A gene of different strains are also shown in supplementary figure S5.

**Figure 5.**
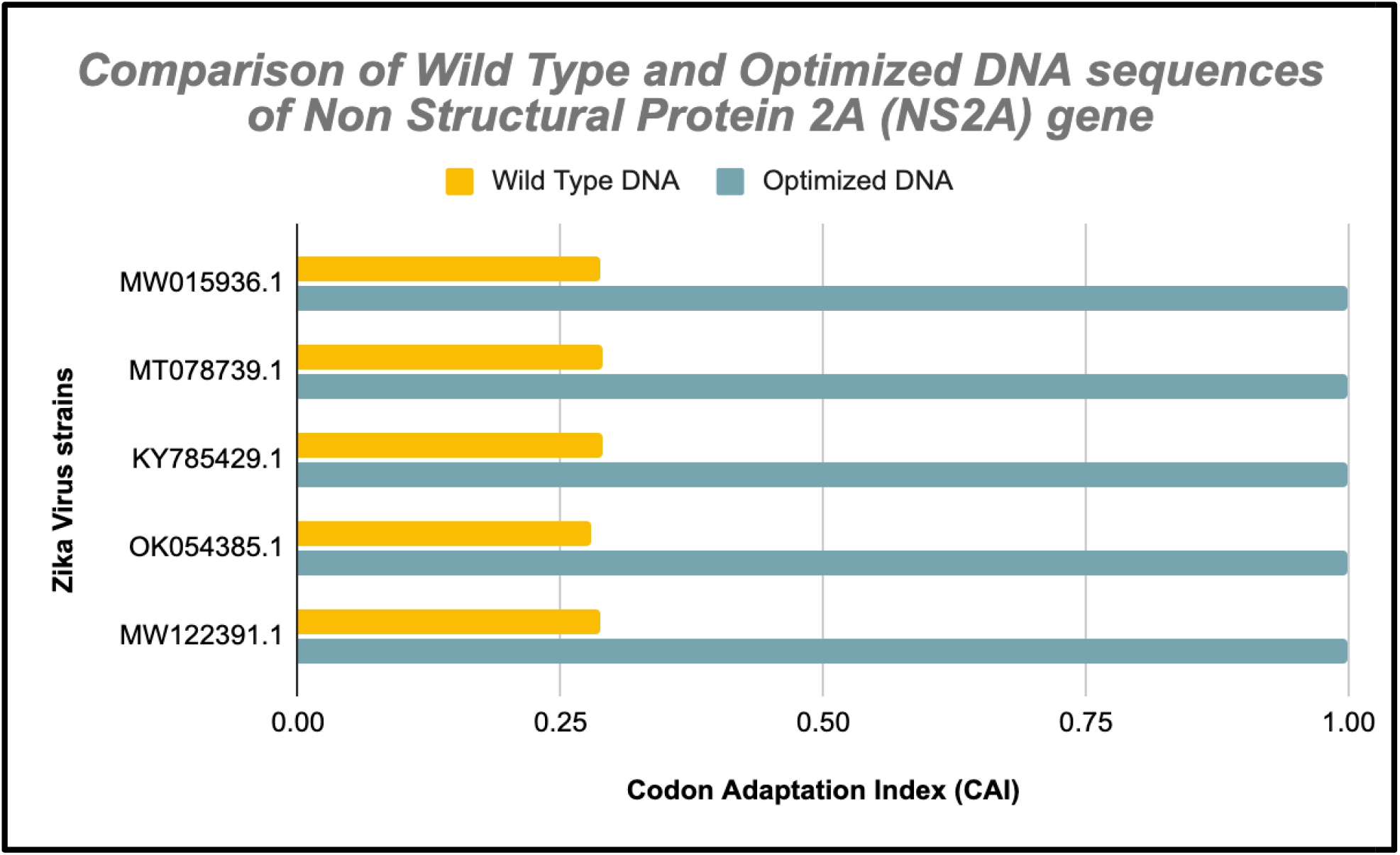
Comparison of the wild-type and optimized DNA sequences of non-structural protein 2A (NS2A) gene in different strains of ZIKA virus.

Then, the CAI, GC% and AT% frequencies in strains of wild-type non-structural protein 2B (NS2B) range from 0.234 to 0.239, 52.6 to 53.6 and 46.4 to 47.4 respectively with an average (±SD) of 0.251(±0.009), 53.3(±0.4) and 46.6(±0.4) respectively. Their respective frequencies in optimized DNA range from 1 to 1, 55.4 to 55.6 and 44.4 to 46.4 with an average (±SD) of 1(±0), 55.4(±0.08) and 44.5(±0.08) respectively (Table 7). The mean CAI and GC in optimized DNA were found to be 3.98 (298.4 %) and 1.03 (3.9%) fold higher than respective values of wild-type. Though, the mean of AT content in optimized DNA was decreased by 4.5% compared to wild-type.

**Table 7.** Non-structural protein 2B gene of Zika virus expression level in *E. coli* of the wild- type and optimized DNA sequences.

| Zika Virus Strain<br>(GenBank<br>Accession<br>number) | Wild-type DNA | Optimized DNA |
| --- | --- | --- |

|  | CAI | GC% | AT% | CAI | GC% | AT% |
| --- | --- | --- | --- | --- | --- | --- |
| MW015936.1 | 0.234 | 52.6 | 47.4 | 1 | 55.6 | 44.4 |
| MT078739.1 | 0.255 | 53.6 | 46.4 | 1 | 55.4 | 44.6 |
| KY785429.1 | 0.255 | 53.6 | 46.4 | 1 | 55.4 | 44.6 |
| OK054385.1 | 0.255 | 53.6 | 46.4 | 1 | 55.4 | 44.6 |
| MW122391.1 | 0.259 | 53.3 | 46.7 | 1 | 55.4 | 44.6 |
| N | 5 | 5 | 5 | 5 | 5 | 5 |
| Min. | 0.234 | 52.6 | 46.4 | 1 | 55.4 | 44.4 |
| Max. | 0.259 | 53.6 | 47.4 | 1 | 55.6 | 44.6 |
| Mean ± SD | 0.251 ± 0.009 | 53.3 ± 0.4 | 46.6 ± 0.4 | 1 ± 0 | 55.4 ± 0.08 | 44.5 ± 0.08 |

A graph was plotted and the CAI value was shown at X-axis, while the number of studied strains on Y-axis (Figure 6). The alignment of the wild and optimized sequences for the NS2B gene of different strains are also shown in supplementary figure S6.

**Figure 6.**
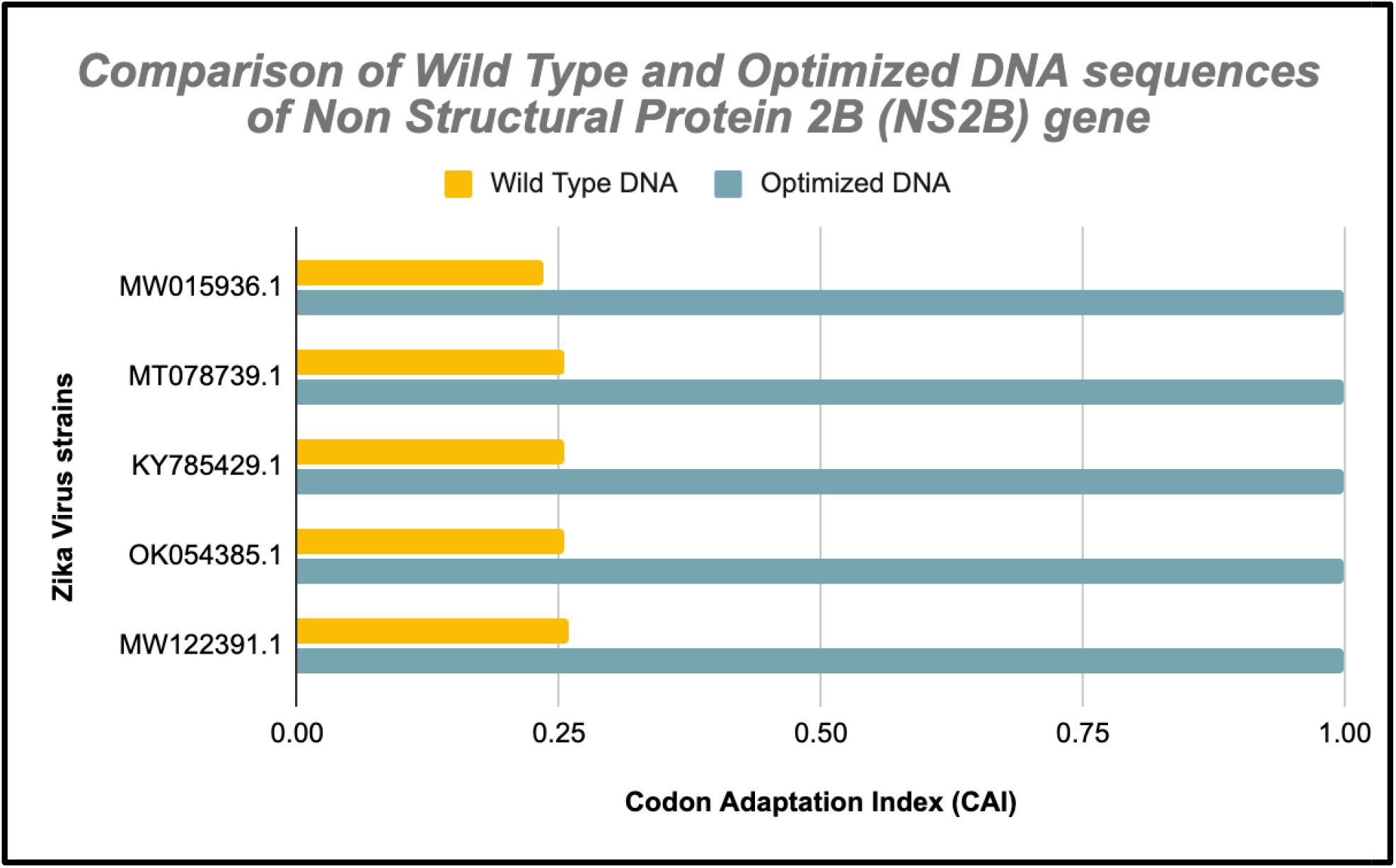
Comparison of the wild-type and optimized DNA sequences of non-structural protein 2B (NS2B) gene in different strains of ZIKA virus.

Furthermore, the CAI, GC% and AT% frequencies of Non-structural protein 3 (NS3) in wild-type strains range from 0.241 to 0.269, 46.5 to 51.0 and 49.0 to 53.5 respectively with an average (±SD) of 0.249(±0.01), 49.3(±2.24) and 50.6(±2.24) respectively. The respective frequencies of these in optimized DNA range from 1 to 1, 51.2 to 55.9 and 44.1 to 48.8 with an average (±SD) of 1(±0), 54.5(±2.0) and 45.4(±2.0) (Table 8). The mean CAI and GC in optimized DNA were found to be 4.0 (301.6%) and 1.1 (10.5%) fold higher than respective wild-type strains. But, the mean of AT content in optimized DNA was decreased by 10.2% compared to wild-type.

**Table 8.** Non-structural protein 3 (NS3) gene of Zika virus expression level in *E. coli* of the wild- type and optimized DNA sequences.

| Zika Virus Strain<br>(GenBank Accession<br>number) | Wild-type DNA |  |  | Optimized DNA |  |  |
| --- | --- | --- | --- | --- | --- | --- |
|  | CAI | GC% | AT% | CAI | GC% | AT% |
| MW015936.1 | 0.243 | 50.9 | 49.1 | 1 | 55.8 | 44.2 |
| MT078739.1 | 0.241 | 51.0 | 49.0 | 1 | 55.8 | 44.2 |
| KY785429.1 | 0.253 | 46.5 | 53.5 | 1 | 53.9 | 46.1 |
| OK054385.1 | 0.241 | 51.0 | 49.0 | 1 | 55.9 | 44.1 |
| MW122391.1 | 0.269 | 47.3 | 52.7 | 1 | 51.2 | 48.8 |
| N | 5 | 5 | 5 | 5 | 5 | 5 |
| Min. | 0.241 | 46.5 | 49.0 | 1 | 51.2 | 44.1 |
| Max. | 0.269 | 51.0 | 53.5 | 1 | 55.9 | 48.8 |
| Mean ± SD | 0.249 ± 0.01 | 49.3 ± 2.24 | 50.6 ± 2.24 | 1 ± 0 | 54.5 ± 2.0 | 45.4 ± 2.0 |

A graph was plotted and the CAI value was shown at X-axis, while the number of studied strains on Y-axis (Figure 7). The alignment of the wild and optimized sequences for the NS3 gene of different strains are also shown in supplementary figure S7.

**Figure 7.**
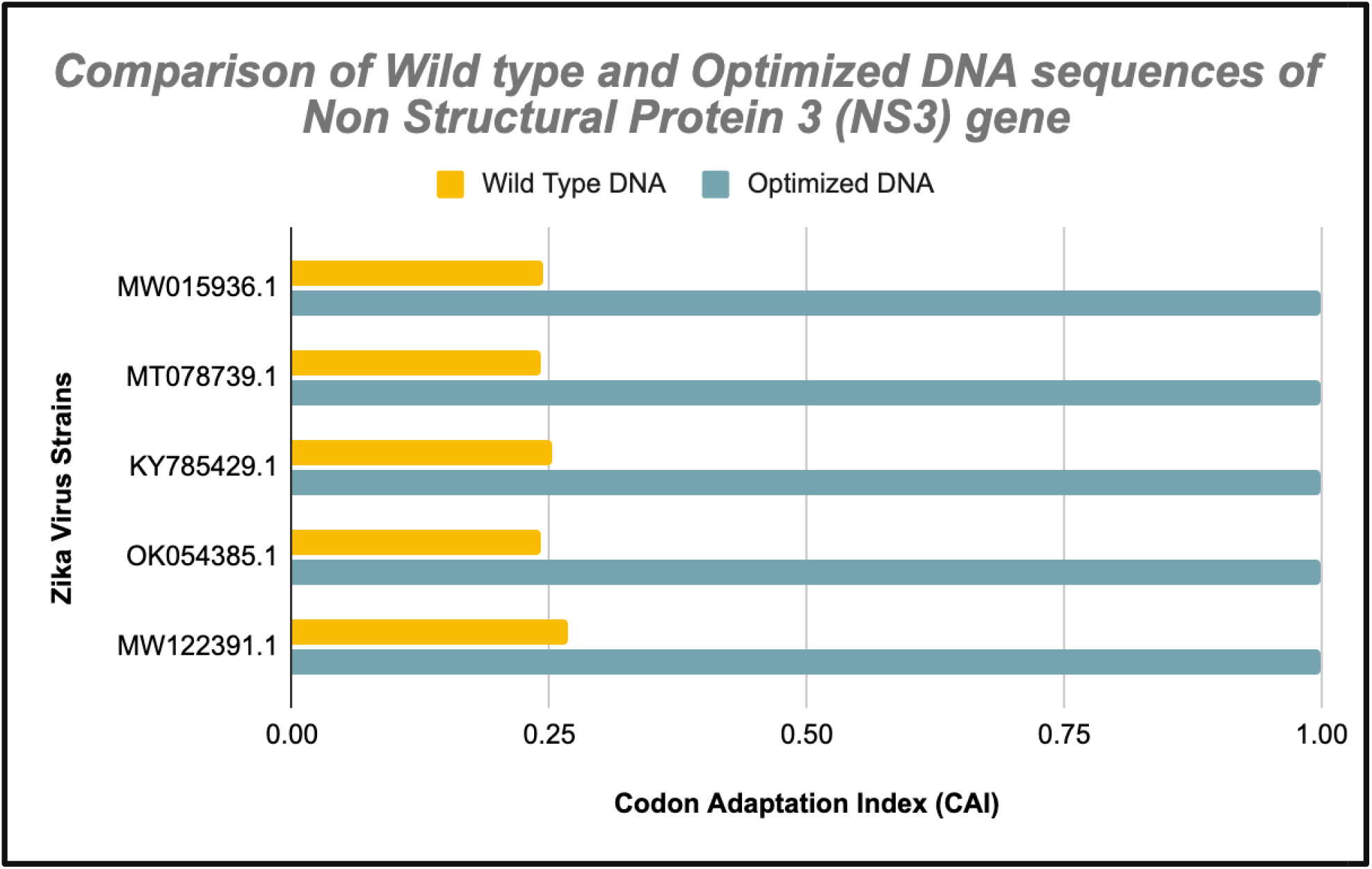
Comparison of the wild-type and optimized DNA sequences of non-structural protein 3 (NS3) gene in different strains of ZIKA virus.

The CAI, GC% and AT% frequencies in six strains of wild-type Non-structural protein 4A (NS4A) ranged from 0.211 to 0.218, 53.8 to 54.9 and 45.1 to 46.2 respectively with an average (±SD) of 0.215(±0.003), 54.02(±0.49) and 45.98(±0.49) respectively. The respective frequencie of these in optimized DNA range from 1 to 1, 57 to 57 and 43 to 43 with an average (±SD) of 1(±0), 57(±0) and 43(±0) respectively (Table 9). The mean CAI and GC in optimized DNA were 4.65 (365.1%) and 1.05 (5.5%) fold higher than the respective mean values of wild-type. However, the mean of AT content in optimized DNA was decreased by 6.4% compared to the wild-type.

**Table 9.** Non-structural protein 4A (NS4A) gene of Zika virus expression level in *E. coli* of the wild-type and optimized DNA sequences.

| Zika Virus Strain<br>(GenBank<br>Accession number) | Wild-type DNA | Optimized DNA |
| --- | --- | --- |

|  | CAI | GC% | AT% | CAI | GC% | AT% |
| --- | --- | --- | --- | --- | --- | --- |
| MW015936.1 | 0.213 | 54.9 | 45.1 | 1 | 57 | 43 |
| MT078739.1 | 0.218 | 53.8 | 46.2 | 1 | 57 | 43 |
| KY785429.1 | 0.218 | 53.8 | 46.2 | 1 | 57 | 43 |
| OK054385.1 | 0.216 | 53.8 | 46.2 | 1 | 57 | 43 |
| MW122391.1 | 0.211 | 53.8 | 46.2 | 1 | 57 | 43 |
| N | 5 | 5 | 5 | 5 | 5 | 5 |
| Min. | 0.211 | 53.8 | 45.1 | 1 | 57 | 43 |
| Max. | 0.218 | 54.9 | 46.2 | 1 | 57 | 43 |
| Mean ± SD | 0.215 ± 0.003 | 54.02 ± 0.49 | 45.98 ± 0.49 | 1 ± 0 | 57.0 ± 0 | 43.0 ± 0 |

A graph was then plotted as described above. The CAI value was shown at X-axis, while the number of studied strains on Y-axis (Figure 8). The alignment of the wild and optimized sequences for the NS4A gene of different strains are also shown in supplementary figure S8.

**Figure 8.**
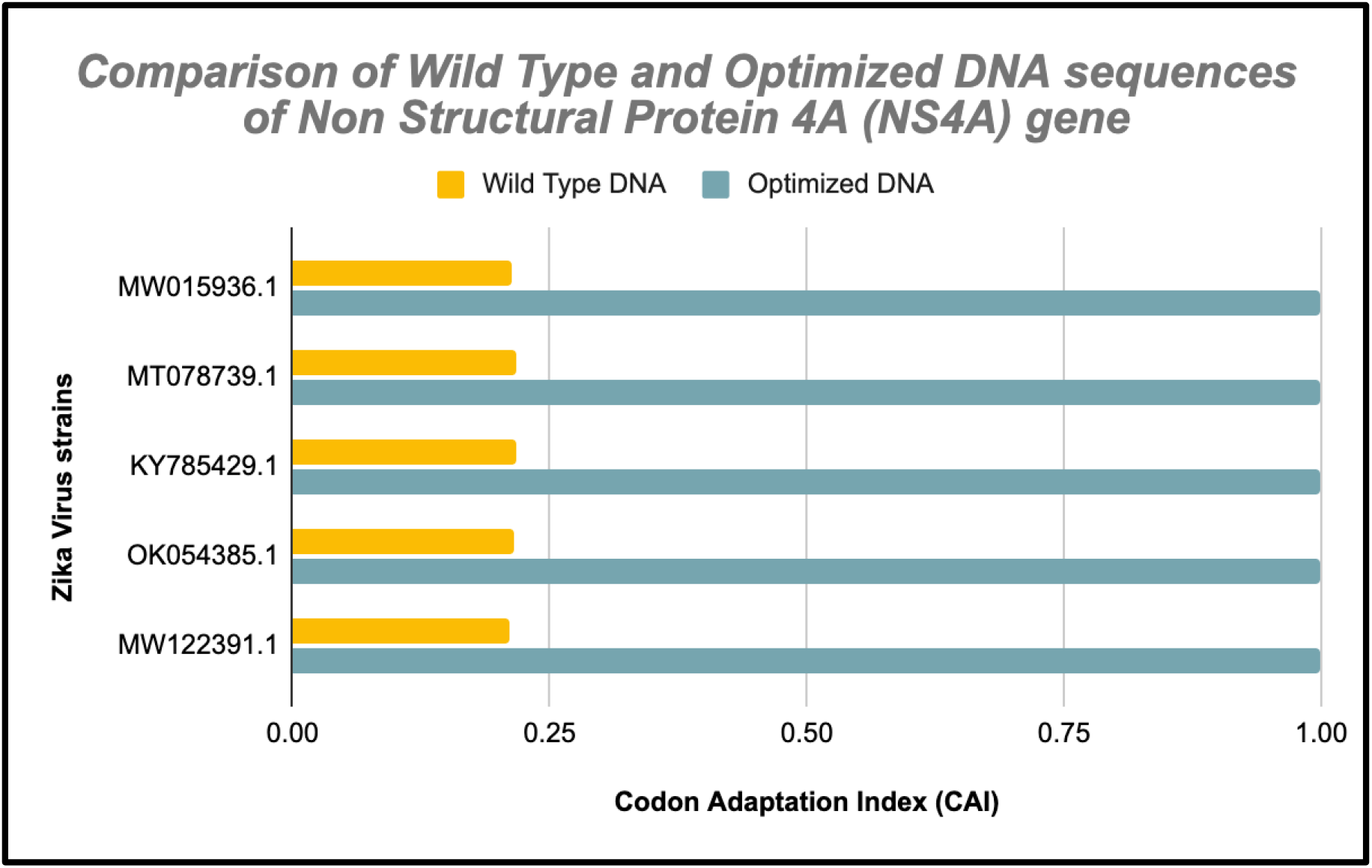
Comparison of the wild-type and optimized DNA sequences of non-structural protein 4A (NS4A) gene in different strains of ZIKA virus.

Similarly, the CAI, GC% and AT% frequencies in strains of wild-type non-structural protein 4 B (NS4B) range from 0.259 to 0.270, 51.8 to 52.6 and 47.4 to 48.2 respectively with an average (±SD) of 0.264(±0.003), 52.26(±0.31) and 47.6(±0.31) respectively. Their respective frequencies in optimized DNA range from 1 to 1, 57.9 to 57.9 and 42.1 to 42.1 with an average (±SD) of 1(±0), 57.9(±0) and 42.1(±0) respectively (Table 10). The mean CAI and GC in optimized DNA were found to be 3.78 (278.7 %) and 1.1 (10.7 %) fold higher than respective values of wild-type. Though, the mean of AT content in optimized DNA was decreased by 11.5 % compared to wild-type.

**Table 10.** Non-structural protein 4 B (NS4B) gene of Zika virus expression level in *E. coli* of the wild-type and optimized DNA sequences.

| Zika Virus Strain<br>(GenBank Accession<br>number) | Wild-type DNA |  |  | Optimized DNA |  |  |
| --- | --- | --- | --- | --- | --- | --- |
|  | CAI | GC% | AT% | CAI | GC% | AT% |
| MW015936.1 | 0.265 | 51.8 | 48.2 | 1 | 57.9 | 42.1 |
| MT078739.1 | 0.270 | 52.6 | 47.4 | 1 | 57.9 | 42.1 |
| KY785429.1 | 0.263 | 52.2 | 47.8 | 1 | 57.9 | 42.1 |
| OK054385.1 | 0.259 | 52.2 | 47.8 | 1 | 57.9 | 42.1 |
| MW122391.1 | 0.264 | 52.5 | 47.5 | 1 | 57.9 | 42.1 |
| N | 5 | 5 | 5 | 5 | 5 | 5 |
| Min. | 0.259 | 51.8 | 47.4 | 1 | 57.9 | 42.1 |
| Max. | 0.270 | 52.6 | 48.2 | 1 | 57.9 | 42.1 |
| Mean ± SD | 0.264 ± 0.003 | 52.26 ± 0.31 | 47.6 ± 0.31 | 1 ± 0 | 57.9 ± 0 | 42.1 ± 0 |

A graph was then plotted for the same. The CAI value was shown at X-axis, while the number of studied strains on Y-axis (Figure 9). The alignment of the wild and optimized sequences for the NS4B gene of different strains are also shown in supplementary figure S9.

**Figure 9.**
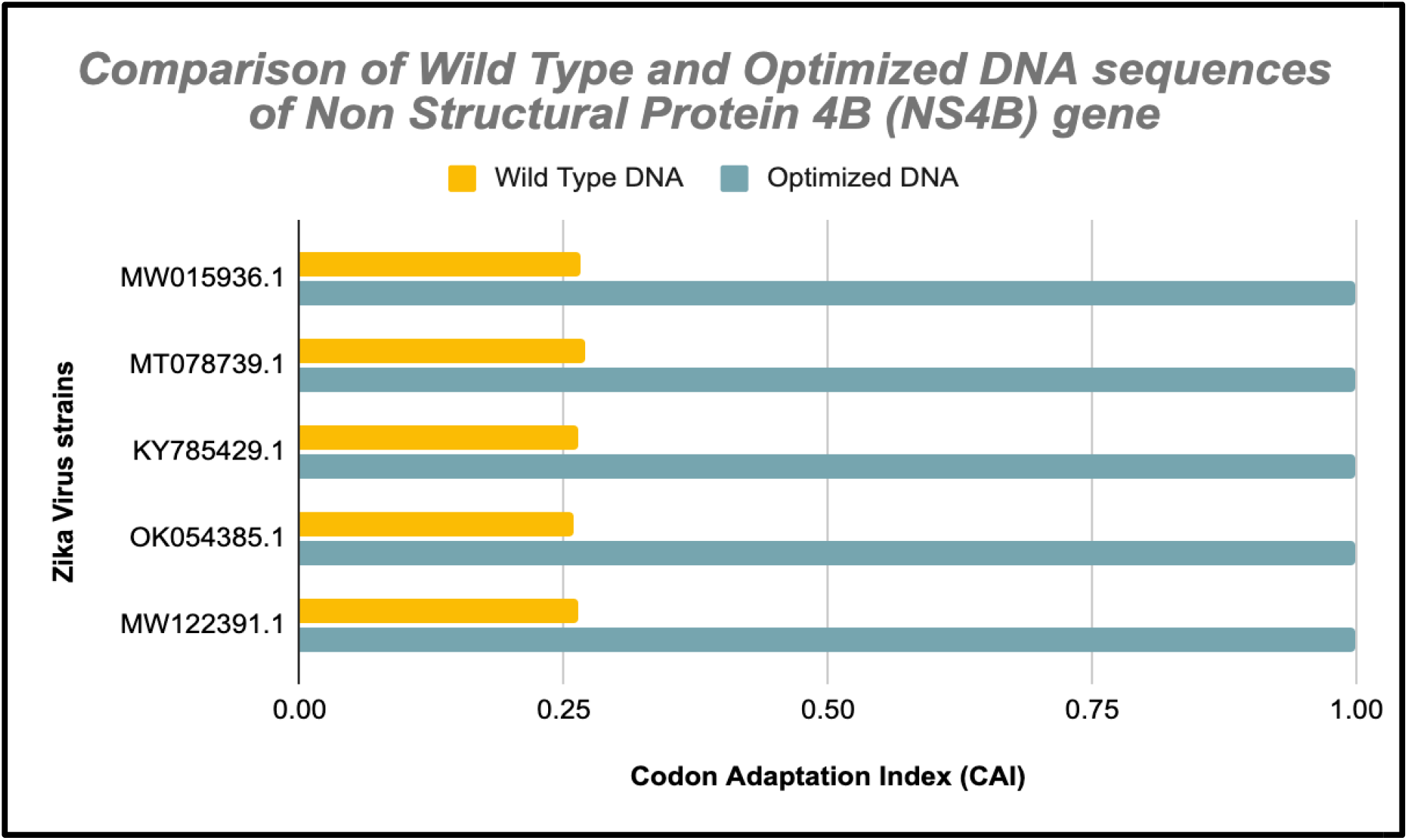
Comparison of the wild-type and optimized DNA sequences of non-structural protein 4 B (NS4B) gene in different strains of ZIKA virus.

Finally, the CAI, GC% and AT% frequencies of Non-structural protein 5 (NS5) in wild-type strains range from 0.267 to 0.272, 51.4 to 54.4 and 45.6 to 48.5 respectively with an average (±SD) of 0.262(±0.015), 52.1(±1.15) and 47.9(±1.15) respectively (Table 11). The respectiv frequencies of these in optimized DNA range from 1 to 1, 53 to 57 and 43 to 47 with an average (±SD) of 1(±0), 54.28(±1.64) and 45.72(±1.64). The mean CAI and GC in optimized DNA were found to be 3.81 (281.6%) and 1.04 (4.18%) fold higher than respective wild-type strains. But, the mean of AT content in optimized DNA was decreased by 4.5 % compared to the wild-type.

**Table 11.** Non-structural protein 5 (NS5) gene of Zika virus expression level in *E. coli* of the wild- type and optimized DNA sequences.

| Zika Virus Strain<br>(GenBank Accession<br>number) | Wild-type DNA |  |  | Optimized DNA |  |  |
| --- | --- | --- | --- | --- | --- | --- |
|  | CAI | GC% | AT% | CAI | GC% | AT% |
| MW015936.1 | 0.267 | 51.5 | 48.5 | 1 | 57 | 43 |
| MT078739.1 | 0.270 | 51.5 | 48.5 | 1 | 53 | 47 |
| KY785429.1 | 0.231 | 54.4 | 45.6 | 1 | 55.4 | 44.6 |
| OK054385.1 | 0.271 | 51.5 | 48.5 | 1 | 53 | 47 |
| MW122391.1 | 0.272 | 51.6 | 48.4 | 1 | 53 | 47 |
| N | 5 | 5 | 5 | 5 | 5 | 5 |
| Min. | 0.267 | 51.4 | 45.6 | 1 | 53 | 43 |
| Max. | 0.272 | 54.4 | 48.5 | 1 | 57 | 47 |
| Mean ± SD | 0.262 ± 0.015 | 52.1 ± 1.15 | 47.9 ± 1.15 | 1 ± 0 | 54.28 ± 1.64 | 45.72 ± 1.64 |

A graph was plotted and the CAI value was shown at X-axis, while the number of studied strain on Y-axis (Figure 10).

**Figure 10.**
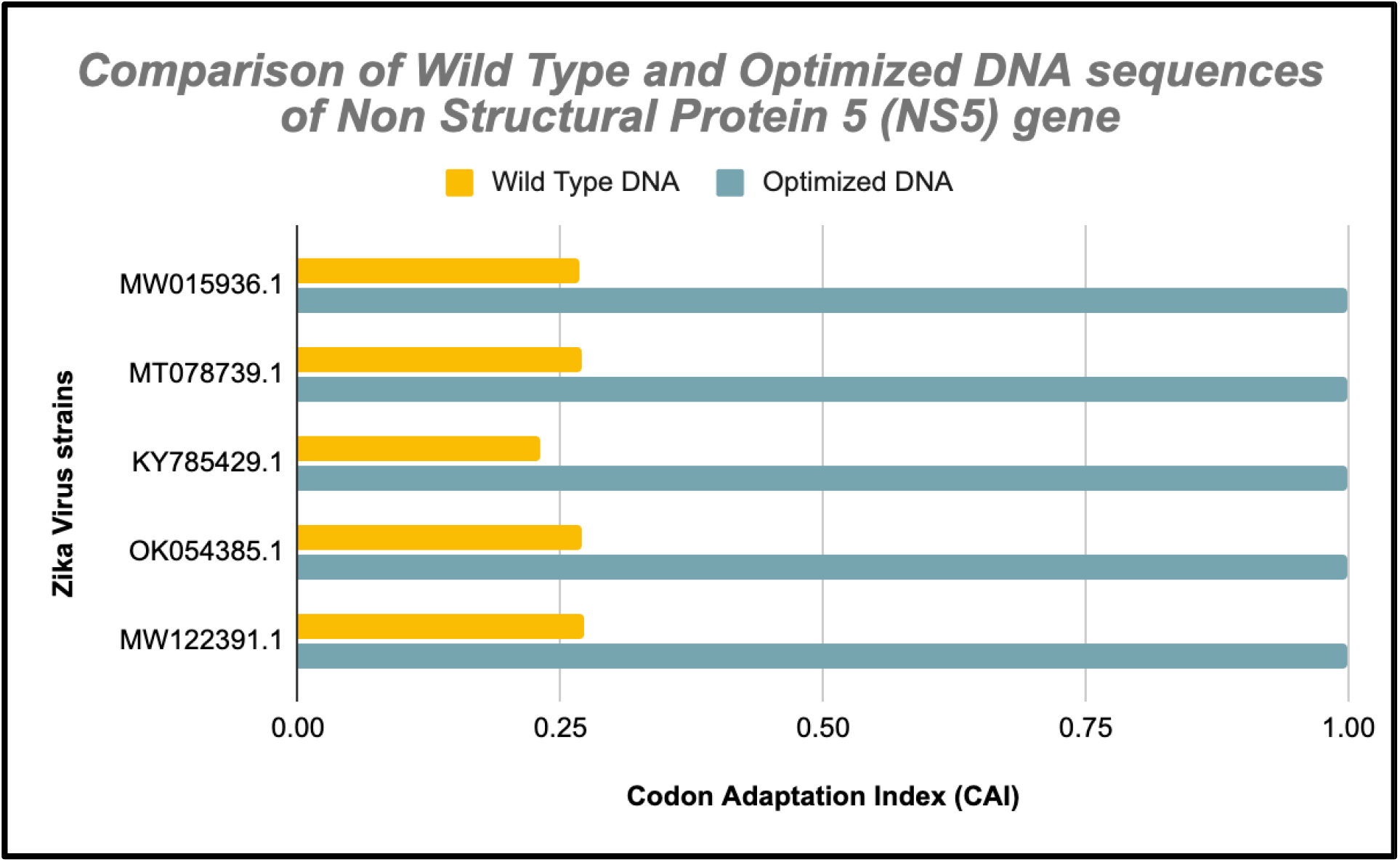
Comparison of the wild-type and optimized DNA sequences of non-structural protein 5 (NS5) gene in different strains of ZIKA virus.

The alignment of the wild and optimized sequences for the NS5 gene of different strains are also shown in supplementary figure S10.

## 4. Discussion

Codon optimization employs the use of high frequency codons to increase protein expression levels which could ensure sufficient and efficient production for research as well as clinical trials, which ultimately results in drug development. There are studies to support the importance of codon optimization like high levels of expression of human iLRP (immature laminin receptor protein), which is a universal tumor antigen, in *E. coli* (Liu et al., 2018); DNA vaccination for HIV (Deml et al., 2001; Lathrop et al., 1999) increasing protein expression levels at lower production costs, removing stop codons, in numerous animal tests etc. The process has many advantages such as matching codon frequencies in host and target for proper folding, inserting/deleting restriction sites, minimizing tandem repeat codons, adjusting rates of translation etc.

In our study we found the average CAI, GC and AT of all strains. The CAI and GC were found to be notably higher in the optimized data than the wild-type while the AT was lower. The CAI of different genes on an average were enhanced by the following values: 4.97 (397.51%) fold for the capsid gene; 3.73 (273.1%) fold for the prM gene; 3.36 (236.7%) fold for the envelope gene; 3.9 (290.6%) fold for the non-structural protein 1; 3.48 (258.4%) fold for the non-structural protein 2A; 3.98 (298.4 %) fold for non-structural protein 2B; 4.0 (301.6%) fold for non-structural protein 3; 4.65 (365.1%) fold for non-structural protein 4A; 3.78 (278.7 %) fold for non-structural protein 4B and 3.81 (281.6%) fold for non-structural protein 5. As compared to wild-type DNA, a relative increase in percentage of GC content was also observed for optimized DNA: 1.04 (4.51%) fold for capsid gene; 1.13(13.9%) fold for prM gene; 1.08(8.46%) fold for envelope gene; 1.05(5.16%) fold for non-structural protein 1; 1.09(9.1%) fold for non-structural protein 2A;1.03 (3.9%) fold for non-structural protein 2B;1.1 (10.5%) fold for non-structural protein 3;1.05 (5.5%) fold for non-structural protein 4A; 1.1 (10.7 %) fold for non-structural protein 4B and 1.04 (4.18%) fold for non-structural protein 5. However, the AT content of genes was lower in all cases. Similarly in silico codon optimization approach has been utilized in various microorganisms such as *Mycobacterium tuberculosis* (Mani et al., 2010), *Aeromonas hydrophila* (Singh et al., 2010), influenza A virus (Mani et al., 20111), SARS-COV- 2 (Al Zamane et al., 2021), Nipah virus (Gupta et al., 2022), etc.

Most ZIKV vaccines currently target precursor membrane protein (prM) as the immunogen. This is a structural protein that is responsible for the assembly of mature virions by cleaving prM to M protein. prM protein has a conserved region in all flaviviruses that is necessary for viral assembly (Yoshii et al., 2012). ZIKV infectivity and pathogenicity can be affected by inhibiting the role of prM during assembly of virus. In most vaccines which are undergoing clinical trials, the structural proteins prM and E are used as antibody activating epitopes (Nambala et al., 2018). The prM/M and E proteins make up the viral envelope of ZIKV and both these are important targets for vaccine design and development. These two structural proteins are determinants for high stability of ZIKV and are non-replicating subunits of the viral genome making them safe candidates for vaccine development. Unlike most flaviviruses which have T-cell epitopes in non-structural proteins, ZIKV has the epitopes for CD4+ and CD8+ T-cell adaptive immunity and neutralizing antibodies in prM and E structural proteins (Chahal et al., 2017; Grifoni et al., 2017; Goo et al., 2018). Most ZIKV vaccines under development are based on plasmid DNA, mRNA platforms or purified inactivated viruses (Tebas et al., 2021; Gaudinski et al., 2018; Modjarrad et al., 2018). These have been unsuccessful when it comes to generating effective antibodies using prM or E proteins. They can, however, be used as a combination of prM-E structural proteins (Nambala et al., 2018). In order to obtain high levels of expression of these genes in cells, codon optimization can be used. When coupled with immunology-based works to test the effective antibodies, effective vaccines can be produced which can be upscaled at industry levels, thus helping us find therapeutics for Zika Virus.

Codon optimization may be the first choice when it comes to generating high gene expressions for therapeutics but it has many limitations as well. There are some serious effects associated with codon optimization which include: optimization may disrupt the normal tRNA patterns affecting the structure and function of proteins; it may reduce efficacy of biotherapeutics; it may lead to production of different peptides with biological activities of unknown origins; post transcriptional modifications may be altered which may lead to modification of protein ensembles. mRNAs have many layers of information overlapping code of amino acids the complexity of which could be disturbed by codon optimization (Mauro et al., 2014) Thus, while codon optimization seems an appropriate choice, one must keep in mind the demerits of the same and analysis for better results is required.

## 5. Conclusions

Zika virus is responsible mainly for asymptomatic infection but severe neurological complications such as congenital Zika syndrome, Guillain-Barré syndrome may also be caused. Microcephaly in infants is also seen. It is primarily spread via mosquitoes. There are no approved vaccines for the virus but the search is on. As vaccines exist for other flaviviruses, there is hope for the ZIKV vaccine as well. After the coronavirus pandemic, ZIKV too poses a threat as a potential pandemic, thus development of safe vaccine and therapeutics is of dire importance. Through this study, we carried out the optimization of the codons for overexpression of different genes of Zika Virus in *E. coli.* These could be used in the development of biotherapeutics. The optimized DNA for every gene for each of the strains had significantly higher CAI and GC% than the wild-type. The ideal genes to be overexpressed in development of biotherapeutics are the membrane precursor (prM) and envelope (E) genes as the results have shown. After *in-silico* studies, *in vitro* studies can help in better validation of the results to determine the levels of overexpression and checking and testing the potency of the vaccines post development. This can be expanded on an industrial scale as well.

## Acknowledgements

Authors are thankful to the Principal, Gargi College for providing the infrastructural support.

---

**Fig. S1.**
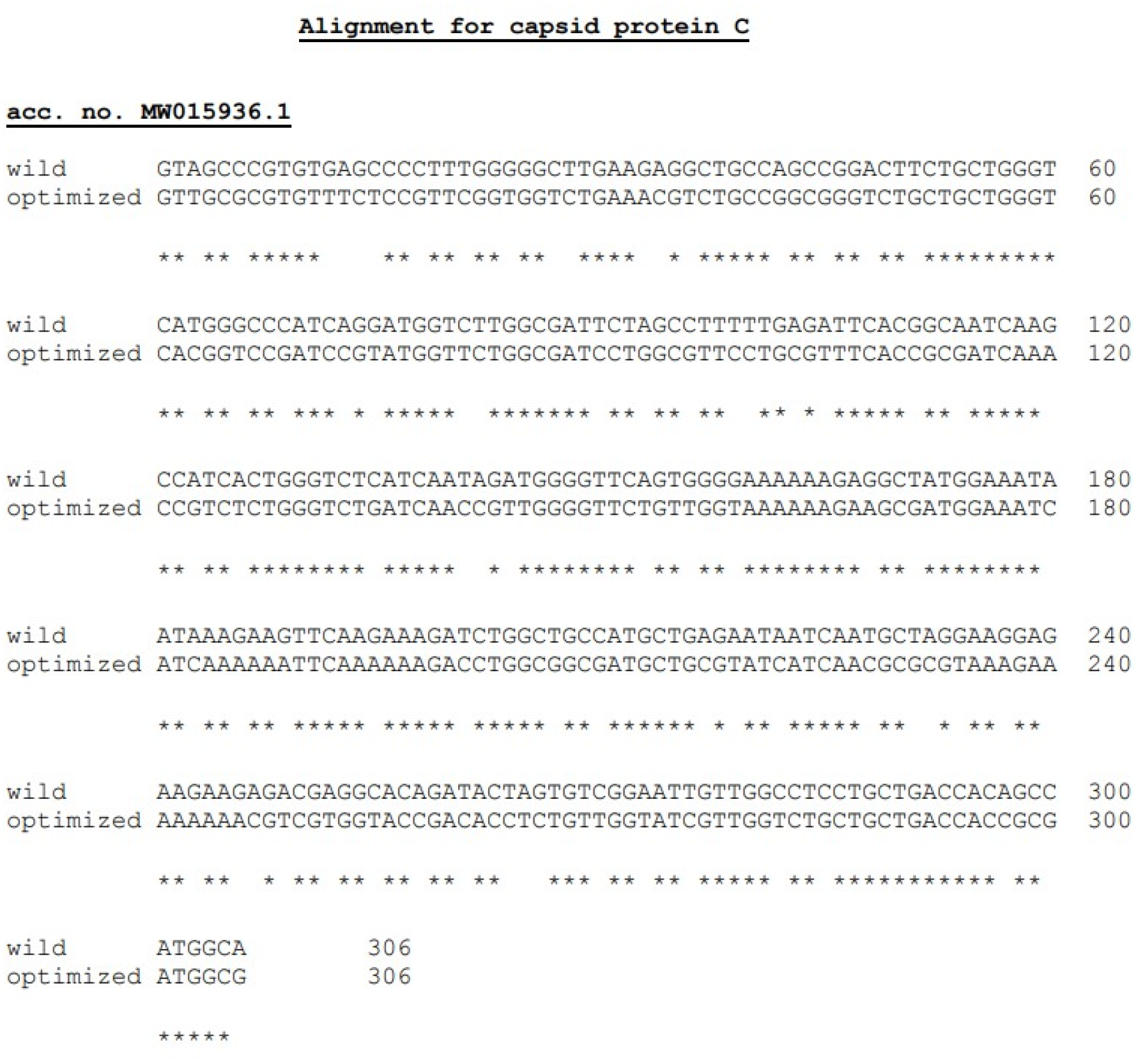

**Fig. S2.**
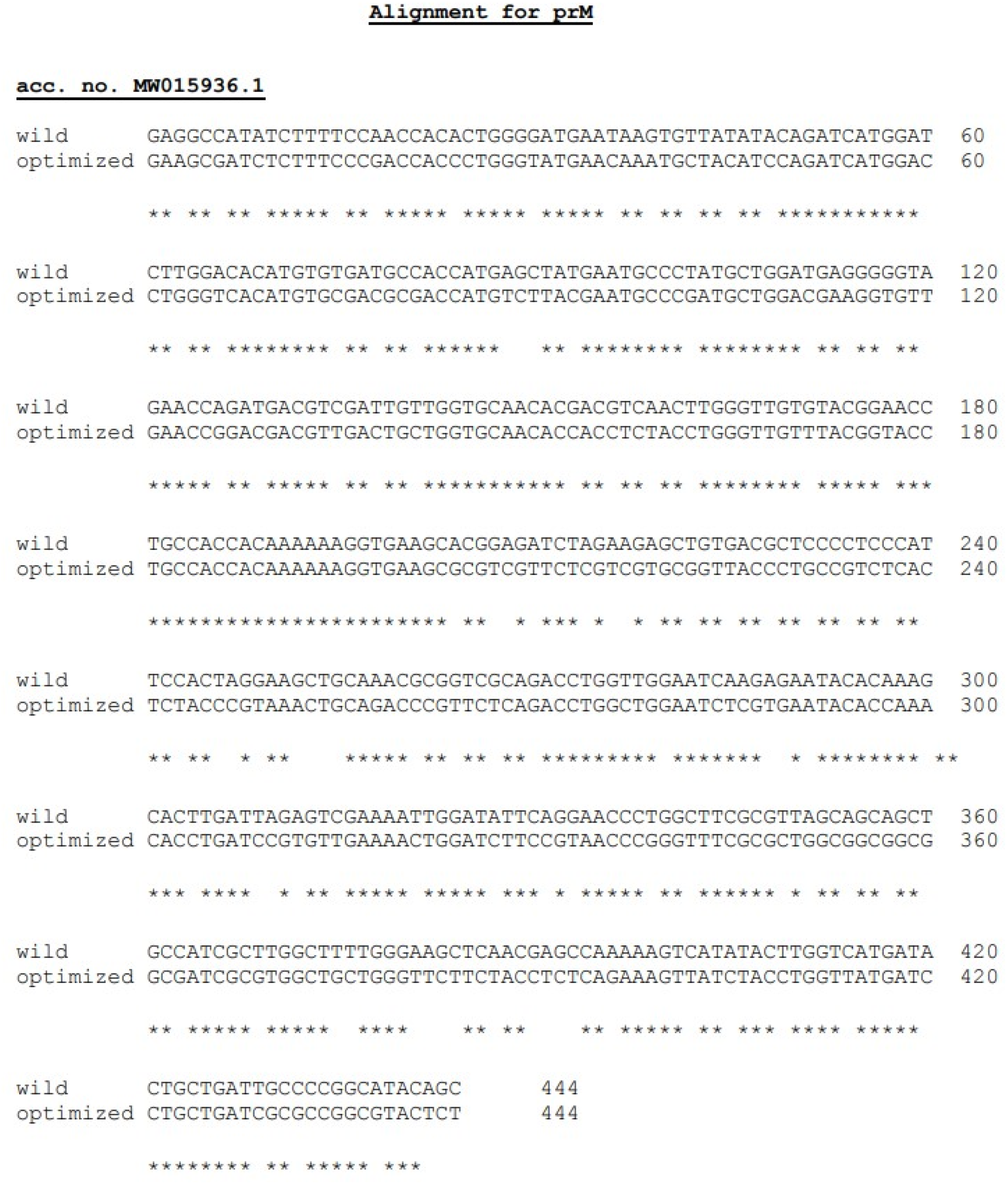

**Fig. S3.**
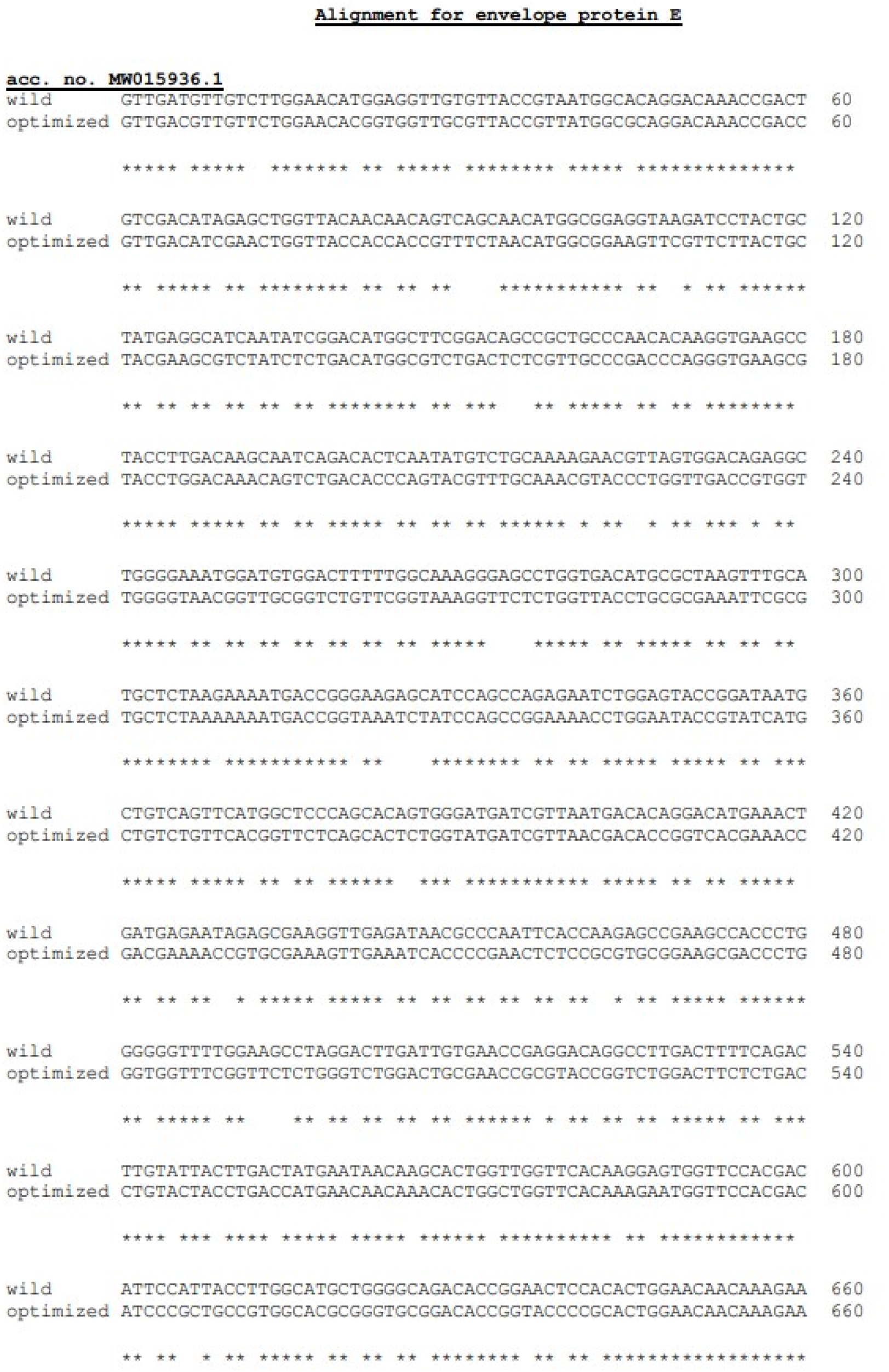

**Fig. S4.**
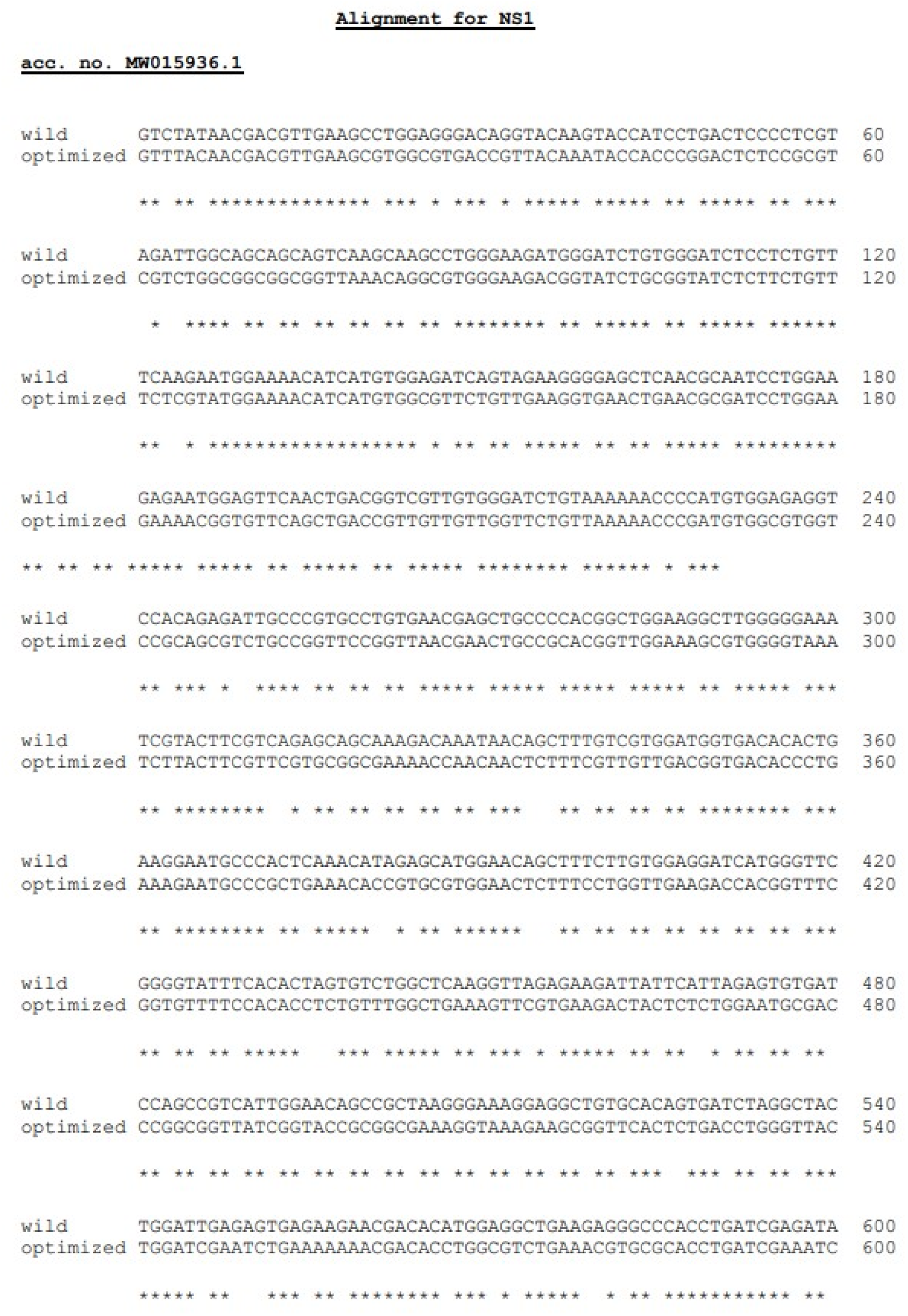

**Fig. S5.**
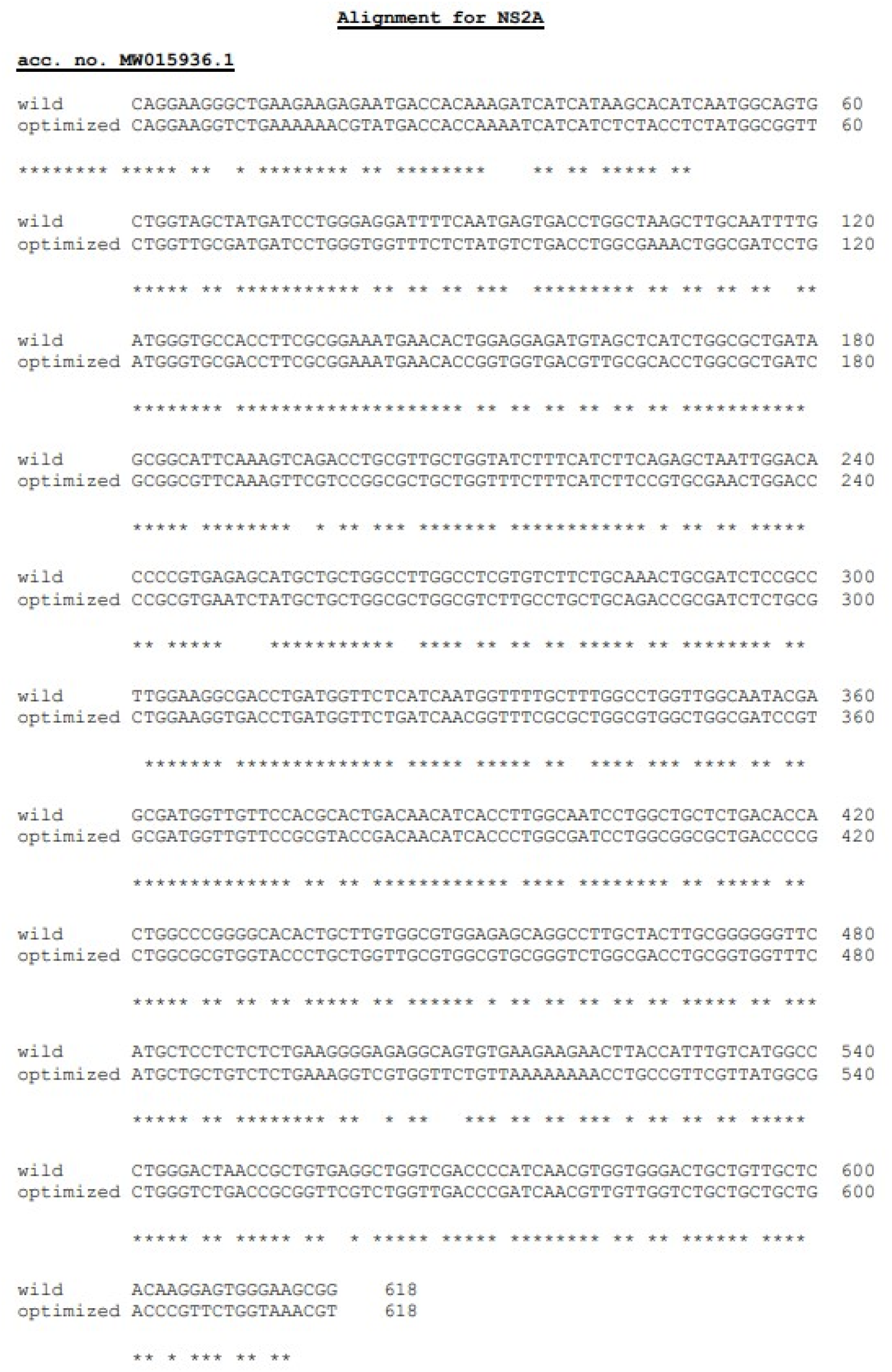

**Fig. S6.**
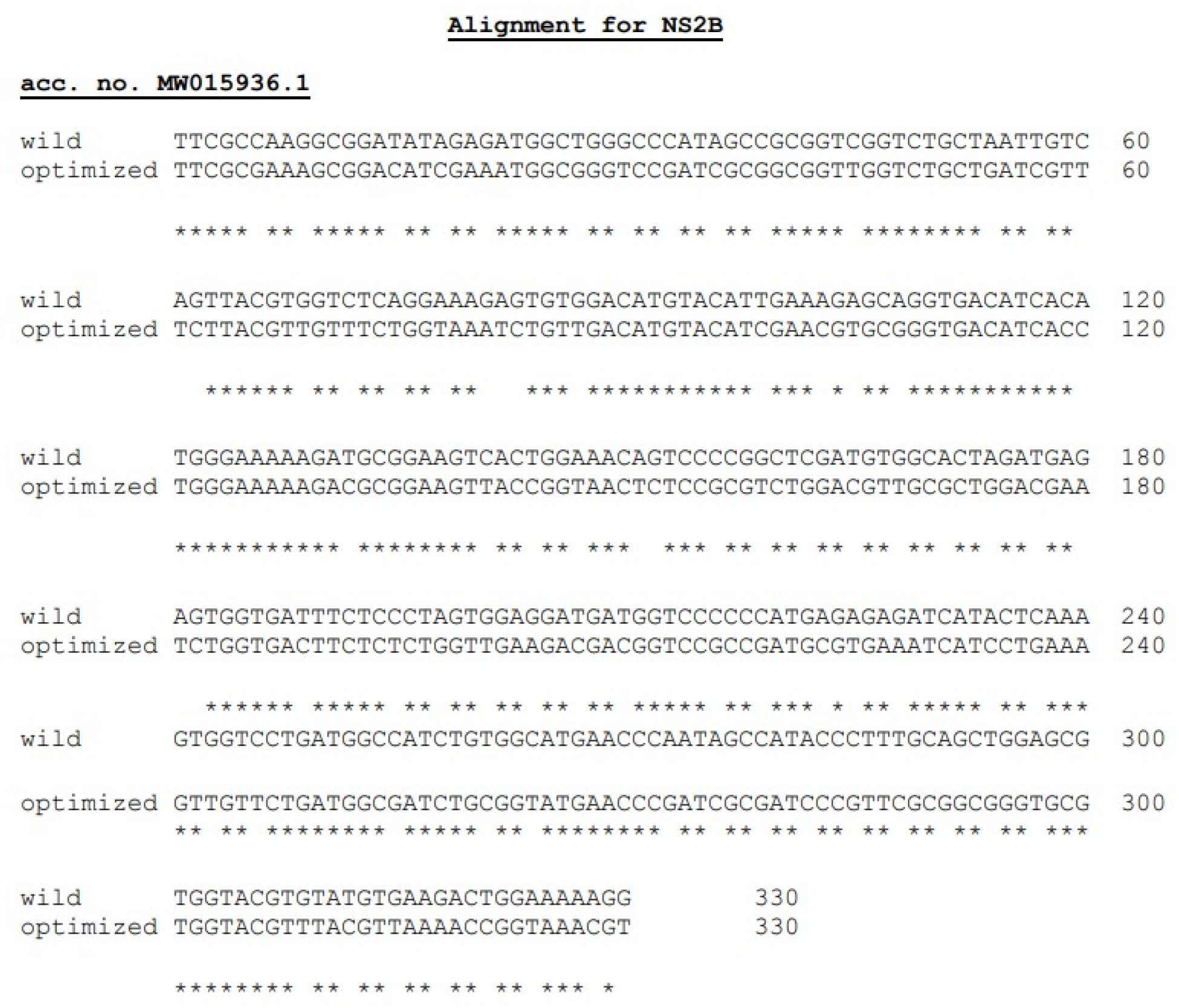

**Fig. S7.**
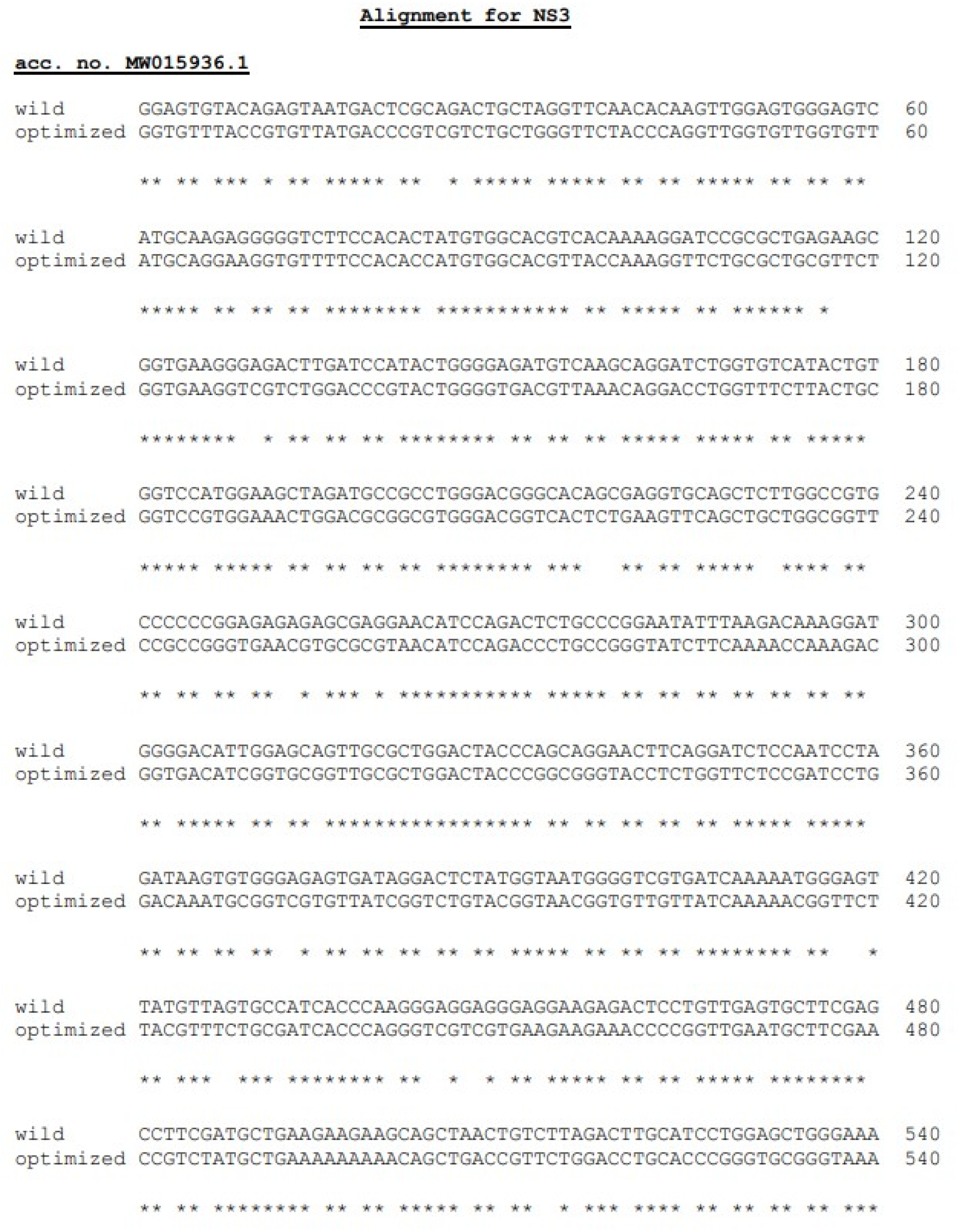

**Fig. S8.**
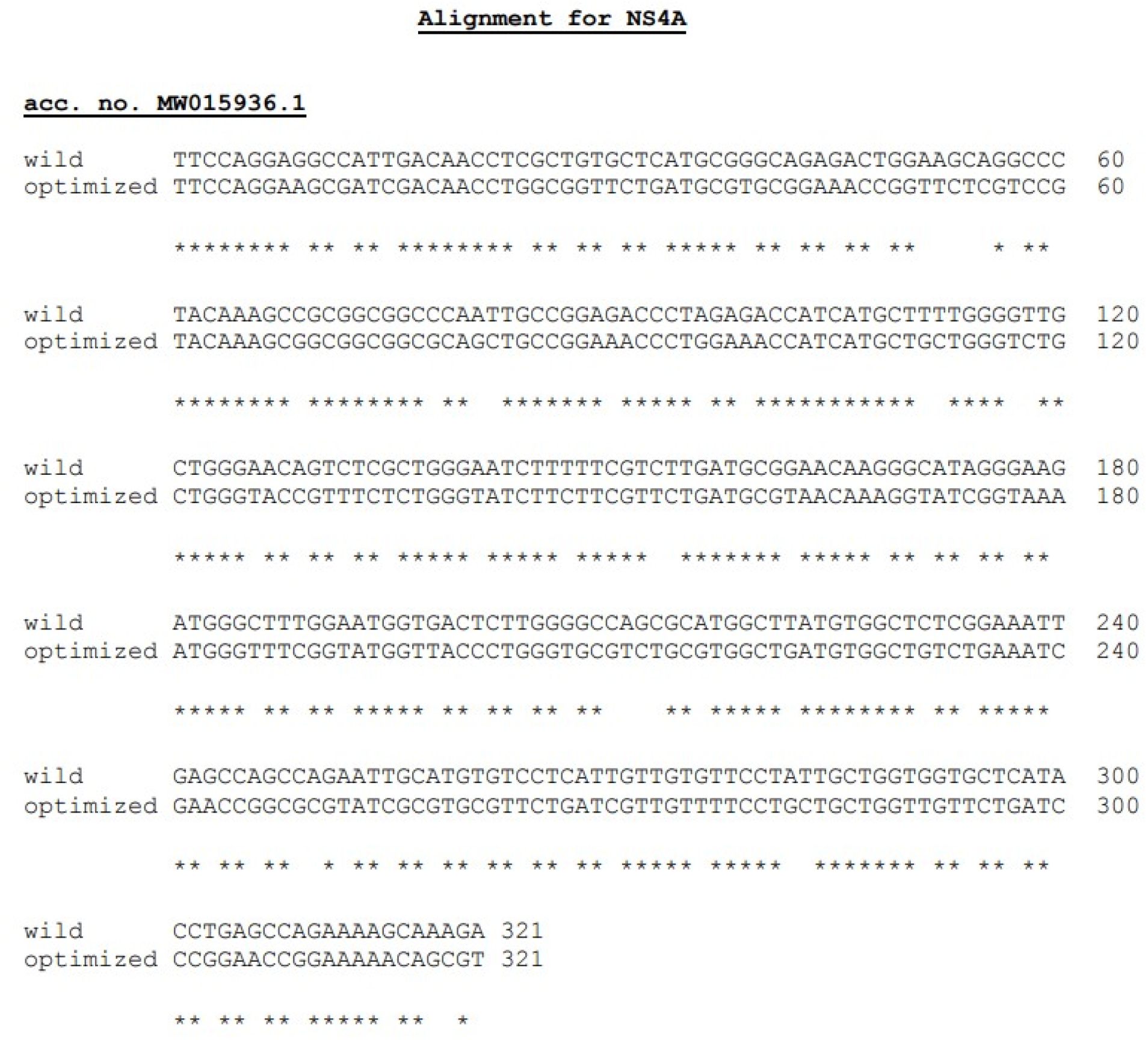

**Fig. S9.**
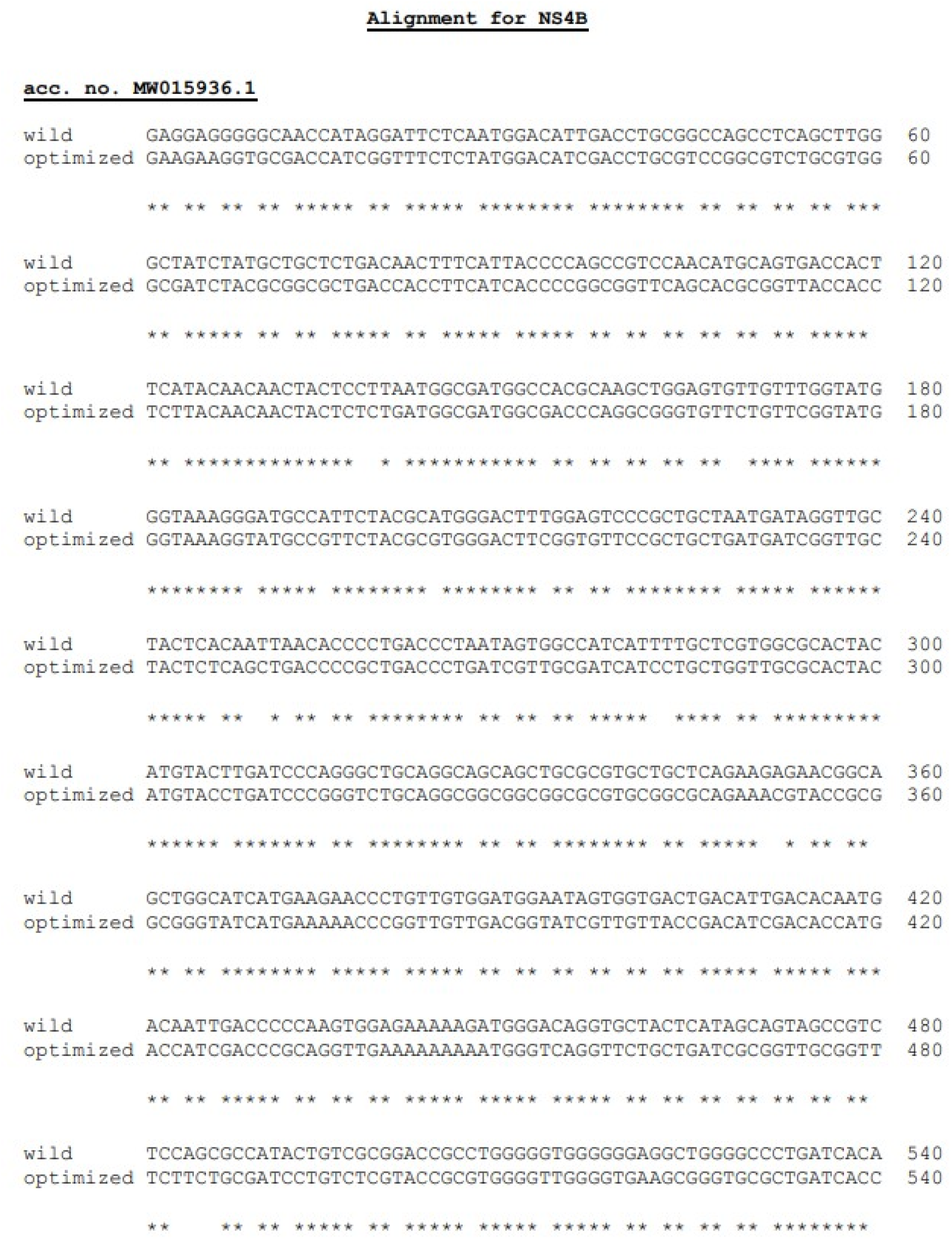

**Fig. S10.**
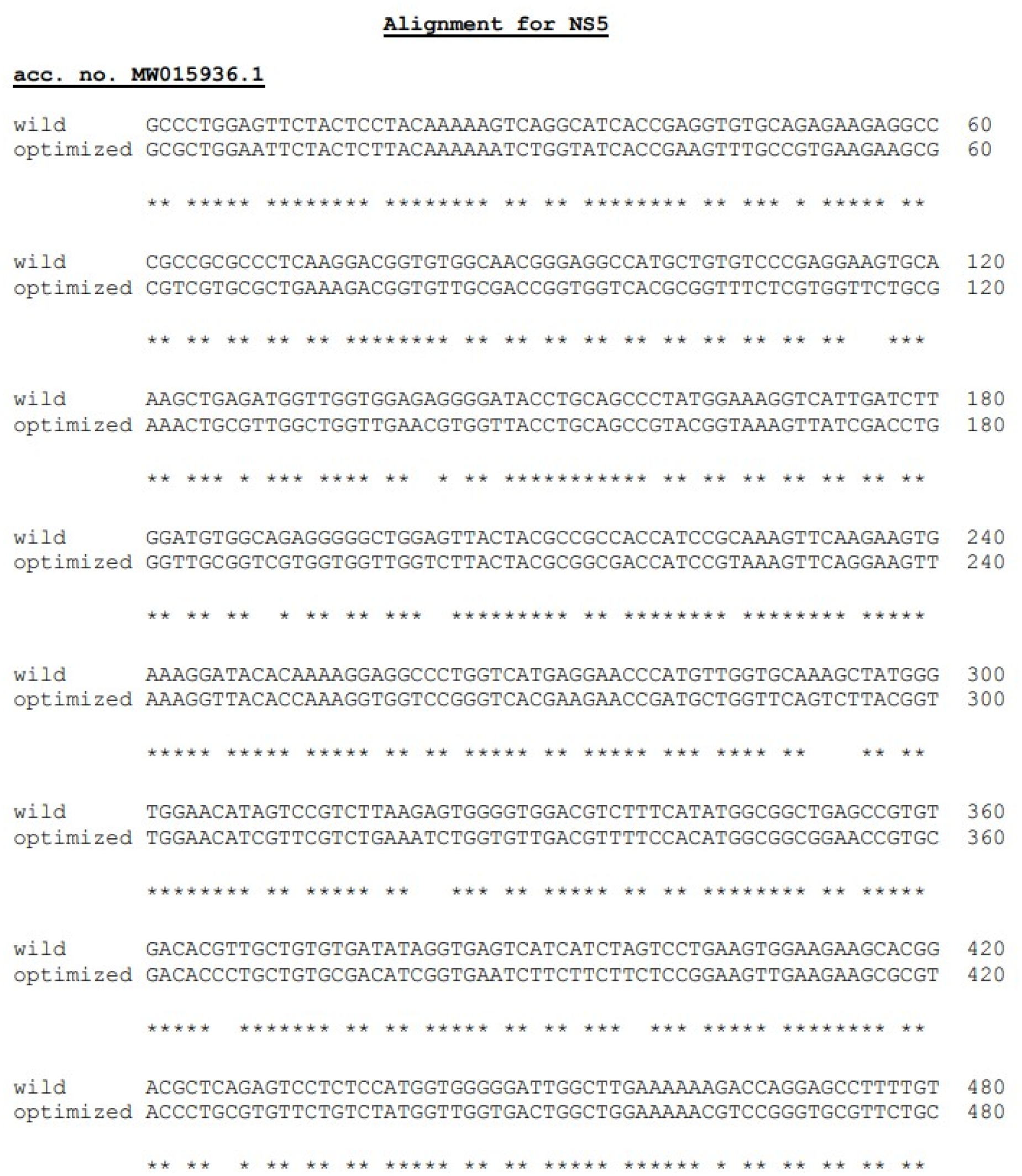

